# SynQuant: An Automatic Tool to Quantify Synapses from Microscopy Images

**DOI:** 10.1101/538769

**Authors:** Yizhi Wang, Congchao Wang, Petter Ranefall, Gerard Joey Broussard, Yinxue Wang, Guilai Shi, Boyu Lyu, Yue Wang, Lin Tian, Guoqiang Yu

## Abstract

**Motivation:** Synapses are essential to neural signal transmission. Therefore, quantification of synapses and related neurites from images is vital to gain insights into the underlying pathways of brain functionality and diseases. Despite the wide availability of synaptic punctum imaging data, several issues are impeding satisfactory quantification of these structures by current tools. First, the antibodies used for labeling synapses are not perfectly specific to synapses. These antibodies may exist in neurites or other cell compartments. Second, the brightness for different neurites and synaptic puncta is heterogeneous due to the variation of antibody concentration and synapse-intrinsic differences. Third, images often have low signal to noise ratio due to constraints of experiment facilities and availability of sensitive antibodies. These issues make the detection of synapses challenging and necessitates developing a new tool to easily and accurately quantify synapses.

**Results:** We present an automatic probability-principled synapse detection algorithm and integrate it into our synapse quantification tool SynQuant. Derived from the theory of order statistics, our method controls the false discovery rate and improves the power of detecting synapses. SynQuant is unsupervised, works for both 2D and 3D data, and can handle multiple staining channels. Through extensive experiments on one synthetic and three real data sets with ground truth annotation or manual labeling, SynQuant was demonstrated to outperform peer specialized synapse detection tools as well as generic spot detection methods, including 4 unsupervised and 11 variants of 3 supervised methods.

**Availability:** Java source code, Fiji plug-in, and test data available at https://github.com/yu-lab-vt/SynQuant.

## 1 Introduction

The synapse is a critical structure in the nervous system that enables communication and interaction between neurons. Cognitive function hinges on proper wiring of synaptic connections within neural circuitry. With the help of microscopic fluorescence imaging of stained antibodies that co-localize with the underlying synaptic cleft, it becomes possible to measure the properties of synaptic puncta and neurites. This information enables researchers to gain insights into how brains function under normal and abnormal conditions. Therefore, automatic and accurate quantification of synaptic puncta is highly needed in today’s brain research. (Lin and Anthony, 2010; Myers, 2012; Ullian et al., 2011; Burette, et al. 2015).

There are two main challenges in analyzing these fluorescence images of synaptic puncta (Fig. 1A, B & C). First, different neurites and puncta show significant variations in terms of morphology and brightness. The reason is the inherent variation among neurons and neurites according to the different roles they play and the discrepancies in maturity. Second, localization of proteins of interest within synaptic puncta is not typically perfect. One possible reason is that there is actually Synapsin I at low concentrations in the neurites, which will show a low level of positive staining. Another possibility is that staining procedures usually result in some degree of “non-specific” staining. This occurs when the antibody containing the fluorophore attaches to something other than the protein of interest. As a result, this diffuse, non-homogenous signal interferes with synaptic punctum detection. For example, even the signal to noise ratio is high for some puncta, it could be much lower for many others in the same data set. The brighter puncta are more likely to be picked up, but this will introduce bias to the analysis. The non-specific antibodies make it hard to identify puncta purely based on intensity. Moreover, some diffused signals could even be stronger than some puncta. Therefore, the combination of punctum-intrinsic heterogeneity, imperfect protein localization to synapses, along with potentially low SNR, lead to great challenges in accurately and reliably detecting, segmenting, and quantifying synaptic puncta.

**Figure 1.**
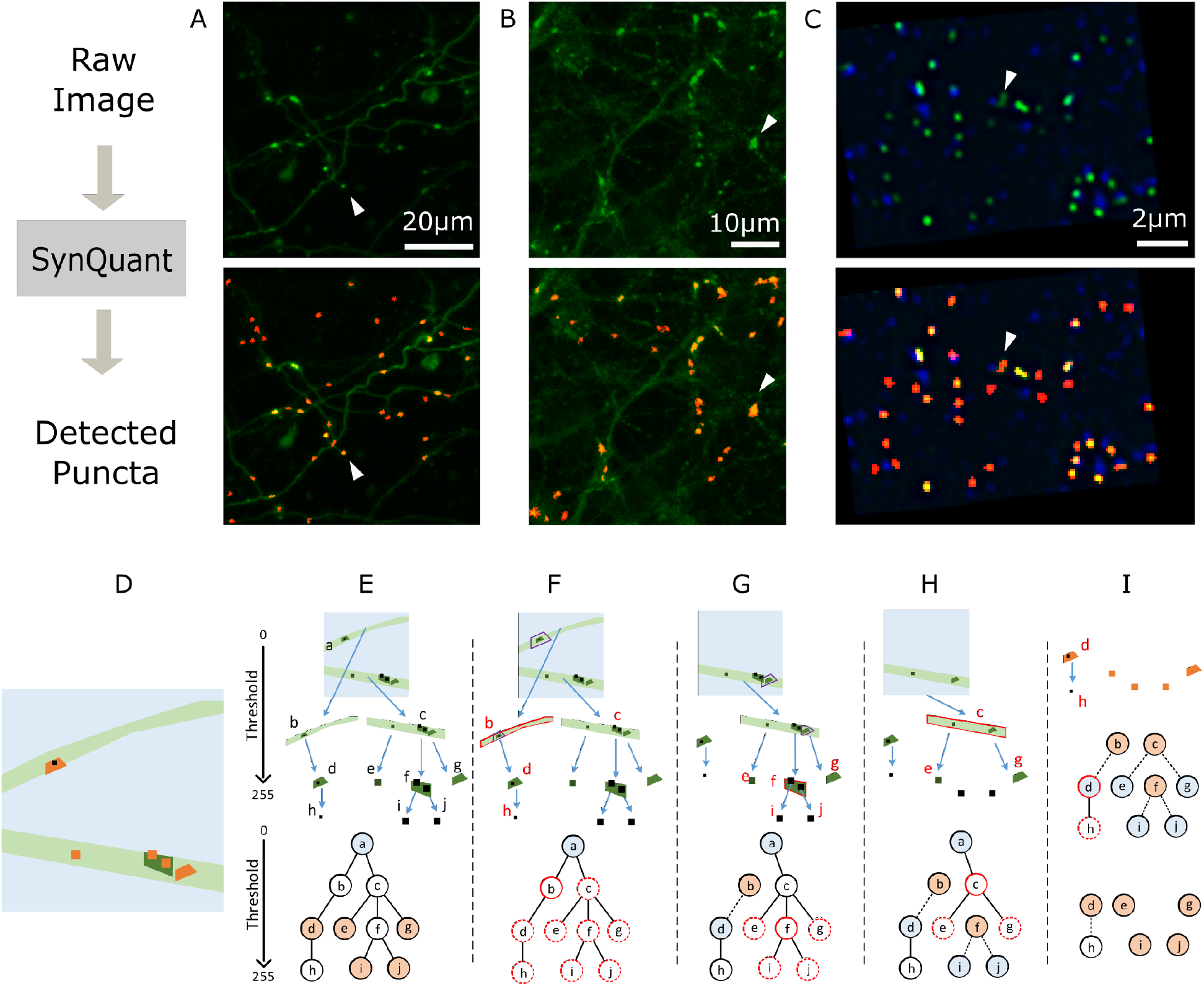
**(A-C)** Examples of raw data and detected puncta. First row images are the raw data and the second row overlaid the detected puncta by SynQuant (shown in red). In the first row, each white arrow points to an example punctum. In the second row, each arrow points to the detected punctum. **(A)** Bass’ 3D in vivo data (mean projected). **(B)** In house neuron-astrocyte co-cultured data. **(C)** Collman’s array tomography data (one z stack is shown). The pre-synaptic channel is shown in blue and the post-synaptic channel is shown in green. The detection results are based on the combination of these two channels. **(D-I)** Joint synaptic punctum detection and segmentation by iterative tree searching and updating. **(D)** Illustration for an image with neurites (light green) and puncta (orange). The light blue background and black dots are both noises from the perspective of synaptic punctum detection. **(E)** Tree structure based on thresholding. Top: the original image is the root node a (*thr=0*). Two branches (b and c) are the children of a with a higher *Thr*. Repeat this process, we get other nodes and edges. Bottom: tree representation. The light blue node is the root and orange ones are the synaptic puncta to be detected. **(F)** b is the current most significant node (red solid circle). The significance of all its descendants d and h, along with all nodes sharing the same ancestry with b are updated (red dashed circles). E.g., the neighborhood of d was originally chosen within a, but now they were chosen within b (purple boxes in a and b). **(G)** d becomes the root of a tree and b is the candidate punctum. As f is the most significant one now, e, i, j and g are chosen to be updated. **(H)** Now we have four trees with a, d, i, and j as roots. Repeat this with node c. **(I)** Continue this process and we get the puncta. There are five significant synaptic puncta detected, d, e, g, i, and j, respectively. Even though b, c, and f are statistically significant regions, they are disqualified as synaptic puncta because they have children that are statistically significant. For the region d, it has a child h, but the region h is not statistically significant, so the region d remains as a synaptic punctum.

Synapse detection has been an active research topic in recent years and quite a few methods were developed (Schmitz et al., 2011; Feng et al., 2012; Danielson and Sang, 2014; Simhal et al., 2017; Simhal et al., 2018; Kulikov et al., 2019). In addition, many image analysis tools for subcellular localization and spot detection have the potential to be repurposed to detect synapses, among which Rezatofighi et al (2012) and Zhang et al (2007) are considered as the state of the art (Smal et al., 2010). We summarized these methods in Table 1 and present their main idea, pros and cons in Table 2. Through experiments on multiple synthetic and real data sets and by comparison to ground truth or human perception, we found the performance of existing algorithms is far from satisfactory, with either high rates of false positives or false negatives. For thresholding-based methods, they do not work well under inhomogeneous background; lack of reliable training data makes it hard to use supervised methods. More importantly, most of them cannot provide a rigorous statistical foundation to assess their output regions and thus give no reliable method to distinguish true synaptic puncta from noises. Besides, the inhomogeneity of synaptic puncta and neurites is not considered and the comparison between images under different conditions is not well calibrated.

**Table 1.**
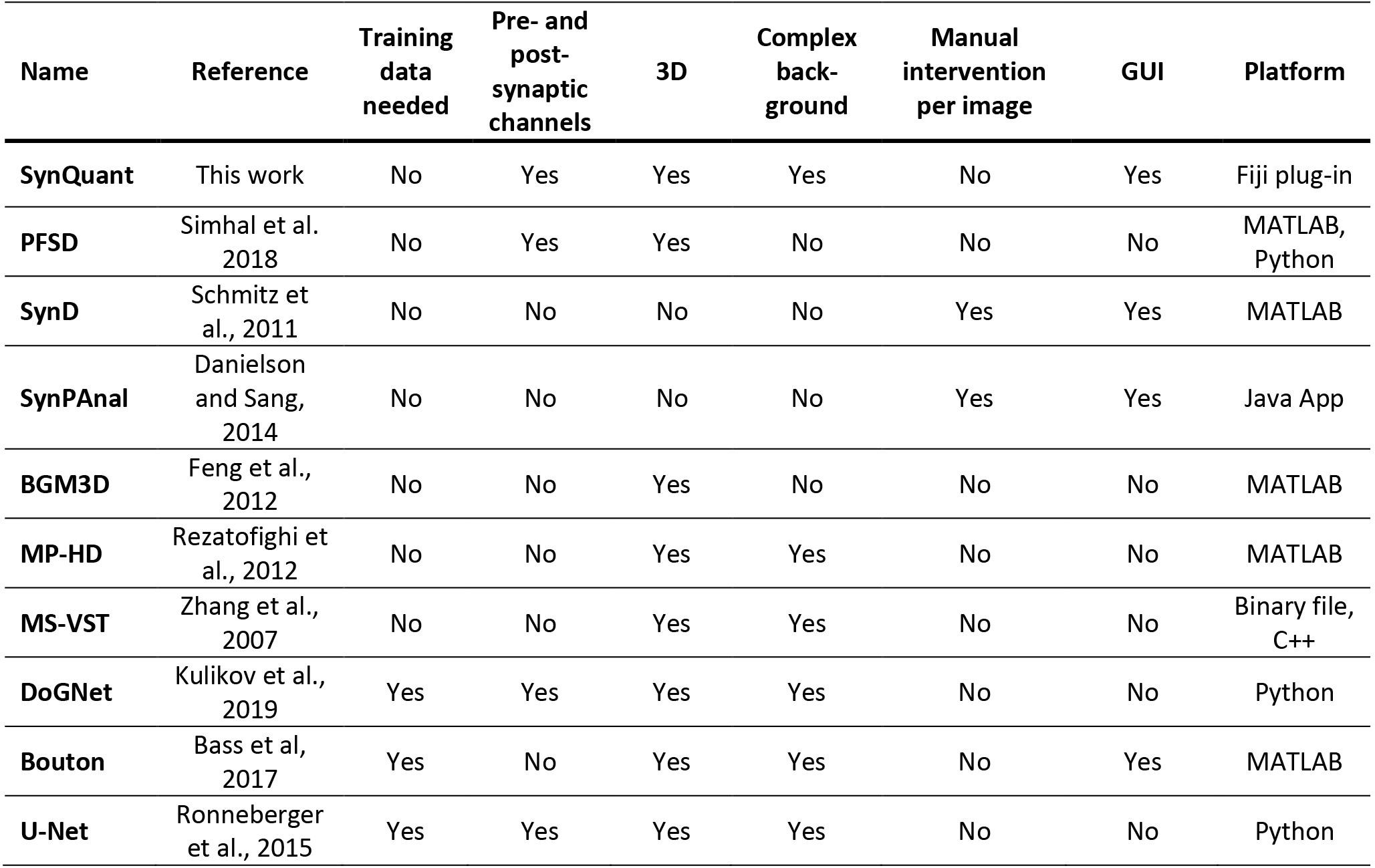
Summary of synaptic punctum and spot detection methods

**Table 2.** Pros and cons of synaptic punctum and spot detection methods

| Name | Detection technique | Pros | Cons |
| --- | --- | --- | --- |
| **PFSD** | Thresholding based on histogram, pixel combining | Simple, scalable and supports multiple channels | Need homogeneous background |
| **SynD** | Template based de-convolution and thresholding | Support soma and dendrite detection, step by step GUI | Need a user specified global threshold per image |
| **SynPAnal** | Thresholding with multiple thresholds | A well-designed GUI | Need manually mark dendrite regions |
| **BGM3D** | Mean-shift based segmentation | Separate overlaid puncta, supports 3D data | Need global thresholding |
| **MP-HD** | Local stable region and H-dome transform | Easy to set parameters | Assume each punctum has a single intensity peak. Consider the intensity contrast only |
| **MS-VST** | Multi-scale wavelet and variance stabilization | Robust in finding weak puncta, directly works with Poisson noise | Do not completely suppress neurite like structures, which lead to false positives |
| **DoGNet** | Artificial neural networks adapted to the shape of synapse | Good detection performance given sufficient training data | Require more training data for deeper models. Less accurate for segmentation unless with pixel level annotation |
| **Bouton** | Interest point detection, local feature extraction and SVM training | Consider local patch information for training | Location of synapse less accurate, cannot segment synapse |
| **U-Net** | Artificial neural networks adapted to biomedical image segmentation | Trainable with relatively small amount of training data | Convolutional kernel not designed for punctum detection |

In this work, we develop a probability-principled synaptic punctum detection method that considers the signal non-specificity, heterogeneity, and large noise. Then we integrate it into our quantification tool (SynQuant) that extracts neurites and synaptic puncta features (Fig. 1 and Supplementary Fig. 1B). To address the signal non-specificity and heterogeneity, we develop a model that is adaptive to localized region properties. If a region is a synaptic punctum, it is expected to be brighter than its surroundings, even though in the same image there may be brighter non-synaptic background regions. Here are two major analytical problems: (1) how to choose the neighborhood pixels for localized modeling and (2) how to evaluate the difference between a candidate region and its surroundings, considering some difference may be purely due to noise. The choice of neighborhood pixels is crucial. For example, for a region inside the neurite, a low intensity pixel in the non-neurite background should not be used as neighbors. A bright pixel in another punctum should not be used either. The difference cannot be solely evaluated based on intensity contrast, because it ignores the number of pixels participating in the comparison: the more pixels, the more reliable the contrast is. Further, although the conventional *t*-test between a group of pixels and their neighbors can integrate the information from intensity contrast and number of pixels, the model is severely biased. The operation of choosing a candidate region and its neighbors has already implied that the candidate region is brighter than its surroundings.

Based on the reasoning above, SynQuant contains two key components. First, we propose to use order statistics (David and Nagaraja, 2003) to properly utilize the local information of puncta and fairly compare all synaptic punctum candidates (Fig. 2). For a given candidate punctum region, SynQuant integrates information from the average intensity inside the region, the average intensity of its neighbors, the sizes of the region and of its neighbors, the ranking of all pixels in these two parts and their noise variance. The theory of order statistics provides a powerful tool to correct the bias introduced by the candidate choosing operation. To the best of our knowledge, this is the first time that the inherent bias for synaptic punctum detection has been rigorously modeled. Indeed, we suspect that the unawareness of the right model for the inherent bias was a major reason for the lack of rigorous statistical model in the field of synapse detection. Second, we propose an iterative updating strategy to identify appropriate neighbors of the synapse candidates for assessing their statistical significance. By this strategy, we will detect the smallest regions retaining statistical significance, which are more likely to be the synaptic puncta. In addition, our method uses the p-value/z-score reported by order statistics to control the false discovery rate, which can be pre-specified by the user. To make the software package comprehensive, we extract neurites by a steerable filter (Meijering et al, 2004) and cut them into roughly homogeneous pieces. For each neurite piece, the position, neurite features, and corresponding synaptic features are computed.

**Figure 2.**
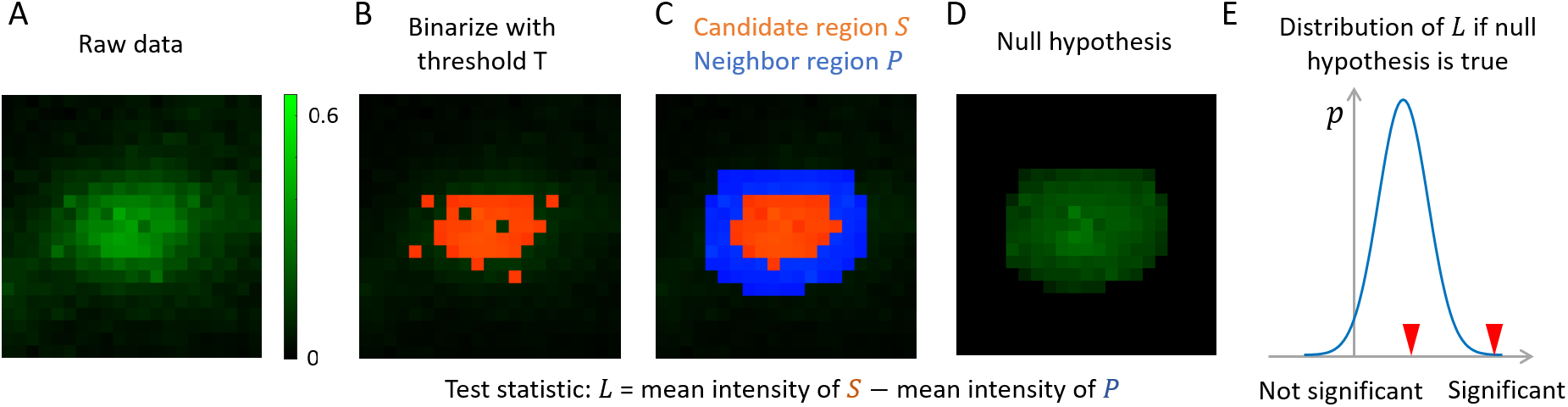
Illustration of order-statistics based punctum significance evaluation. (**A**) A small patch of raw data. Brighter pixels have higher intensities. All pixels are contaminated by Gaussian noise N(0, σ^2^), where σ^2^ is 0.005. (**B**) Binarize (A) with threshold 0.4. The pixels above the threshold is shown in red. (**C**) We remove isolated pixels in (B) and fill small holes. The resulting red part S is a candidate region. The blue pixels form its neighbor region P. We create a sorted list Ω of all pixels in region S ∪ P, where a pixel with higher intensity will appear earlier. In (C), most pixels in S appear earlier than most pixels in P in Ω. (**D**) Under the null hypothesis, there is no true punctum and we obtain Ω (along with S and P) by chance due to noise. In (D), we show an example when all pixels are gaussian noise N(0.27, 0.005) and the intensity order given in Ω is obeyed. The inner part is still brighter than the neighboring part, but the difference is much less obvious since there is no true punctum now. (**E**) Under the null hypothesis, we use order statistics to obtain the null distribution of L(Eq. 2 to Eq.5). The distribution of L has a positive mean, which models the bias of thresholding operation. For the candidate found in (C), we calculate its test statistic and check it against the null distribution of L. If it is in the position of the left red arrow, the punctum is not significant. If it is in the right red arrow, it is significant. Larger intensity difference between S and P larger size of S, and lower noise level will make the candidate more significant and more likely to be chosen. The creation of the null distribution also models the effect of filling small holes and considers the isolated higher intensity pixels in P.

Experiments show that our quantification framework obtains a large accuracy gain of synaptic punctum detection on both simulated data set and three annotated real data sets. In addition, we build a regression model to study the relationship among those piece-level features and brain conditions or disease phenotypes. Since the inhomogeneity is modeled as a confounding factor, the relationships between synaptic punctum density and disease phenotypes are reliably uncovered. In the rest of the paper, we will use synaptic puncta or puncta to refer the signals in fluorescence imaging that we want to detect. We use synapse or synaptic cleft to refer to the manual annotation in the electron microscope.

## 2 Method

We first estimate the noise model parameters and stabilize the noise variance of the image (Supplementary Fig. 1B, left panel). After that, we create candidate punctum regions by binarizing the image with multiple intensity thresholds. These thresholds cover the whole range of signal intensities and do not require user intervention. Each threshold leads to some binary connected components, or regions (Fig. 1E). Clearly, regions can be overlapped. Indeed, we build a tree structure where each region becomes a node. A child node represents that the corresponding region is completely contained in its parent region. Each region is assigned an initial significance using order statistics. We iteratively search for possible synaptic puncta in the tree and update the statistical significance for each candidate based on the search. The determination of a positive punctum is controlled by the user-specified threshold on the significance level.

The synaptic features are extracted after post-processing. The related neurite attributes are collected from the neurite channel after neurite tracing and are cut into segments (Supplementary Fig. 1B, right panel). The features are then used for statistical analysis along with the disease phenotypic information of each image.

### 2.1 Noise estimation and variance stabilization

Application of order statistics theory requires the noise statistics of the pixels in a candidate region and its neighbor. Conventionally, the noise is modeled as following a Gaussian distribution which simplifies subsequent computations. However, the photon detector introduces noise whose variance is linearly dependent on the signal intensity. We apply the noise model proposed by Foi et al. (2008). The variance for pixel (*i,j*) is modelled as

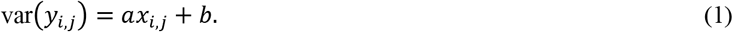

Here var(*y*) is the pixel noise variance. *x* is the underlying signal intensity, which is not observed but can be well approximated by the observed pixel intensity. The term *ax* models the Poisson type noise and the term *b* models the additive Gaussian noise. The model can be fit based on pixel data from a single image and the resulting *a* and *b* are used in the Anscombe transform to stabilize the noise (Foi et al., 2008), so that the noise variance associated with the new values after the transform is independent to the intensity itself and can be approximated by a single constant 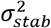.

### 2.2 Synaptic puncta’s significance scores based on order statistics

In our adaptive tree search and updating algorithm, for each threshold, we get a set of isolated regions (nodes in the tree), each containing a set of pixels (Fig. 1D and Fig. 1E). These regions are potential candidates for synaptic puncta that need to be evaluated by statistical tests. The test for the individual region is based on the difference of this region and its neighbor pixels (Fig. 2E). A larger difference implies a larger possibility that this region is significantly different from the surroundings, which is a necessary (but not sufficient) condition for being a synaptic punctum. For each region, a group of neighbor pixels is selected. We assume there are *M* pixels *S* = {*x*_1_, …, *x*_*M*_} in the region and *N* pixels *P* = {*x*_*M*+1_, …, *x*_*M*+*N*_} in the neighbor. *x*_*_s denote the intensity levels of the pixels. We may use a *t*-test to compare these two groups. However, due to the thresholding operation, almost all the *M* pixels have higher intensities than the *N* neighbors, though a few exceptions are allowed like isolated high-intensity pixels in the neighbors or low-intensity holes inside the region (Fig. 2B and Fig. 2C). Even if there is no true signal, due to the thresholding, positive difference usually exists between the means of the two groups for any candidate region considered (Fig. 2D and Fig. 2E). This positive difference is a bias and, if not corrected, will complicate the detection and result in a lot of false detections. Here, we are still interested in the difference between the candidate region and its neighbor pixels, and define the test statistic as the following,

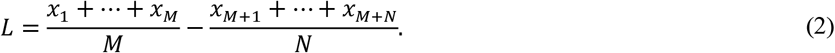

Due to the thresholding, the intensities {*x*_1_, …, *x*_*M*_} are almost always larger than any intensity of {*x*_*M*+1_, …, *x*_*M*+*N*_}, even without a true signal. Thus, *L* will almost always be positive. The exact value of *L* is determined by the noise variance, the sample size and the ratio of *M* and *N*. The theory of order statistics provides a formal approach to account for the bias by calculating the mean and variance of *L* under the null hypothesis that there is no true signal among the candidate region and its neighbor pixels. Let *n* = *M* + *N*, we can rewrite *L* as in (David and Nagaraja, 2003):

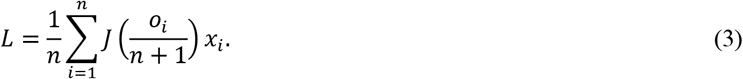

Here, *J*(*k*) is a weight function corresponding to the coefficients for *x*_*i*_ in Eq. (2). For 1 ≤ *i* ≤ *M*, *J*(*o*_*i*_/(*n* + 1)) = *n*/*M*, and for *M* + 1 ≤ *i* ≤ *M* + *N*, *J*(*o*_*i*_/(*n* + 1)) = −*n*/*N*. *o*_*i*_ is the intensity order of *x*_*i*_ among the *n* samples. For instance, *o*_*i*_ =3 if *x*_*i*_ is the pixel with the third highest intensity in the *n* pixels. We write

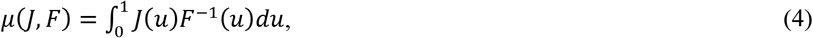

and

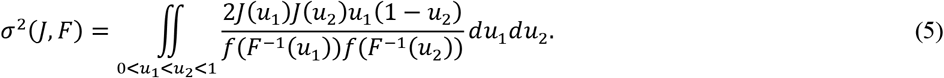

Then we have 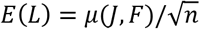 and *var*(*L*) = *σ*^2^(*J*, *F*)/*n*, when *n* = *M* + *N* → ∞ (David and Nagaraja, 2003). Here *f* is the normal probability density function with zero mean and variance as the stabilized noise variance 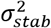. *F*^−1^ is the corresponding inverse normal cumulative distribution function. The integration is computed by summation using all the *n* samples. Then we define the order statistic score *z* as a function *f*_*os*_:

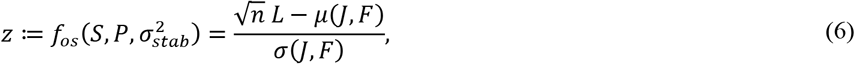

where *z* follows asymptotically a standard Gaussian distribution and hence can be easily used to compute the statistical significance of any observed value of *L*.

We note that the statistical significance computed above is a good approximation only when the sample size is large enough, which may not be the case. With some typical image resolutions, one synaptic punctum may only contain about 10 or fewer pixels. Here we apply two corrections for the small sample size to improve the approximation. First, we notice for the double integration in *σ*^2^(*J, F*), the integration space is a triangle defined by 0 < *u*_1_ < *u*_2_ < 1. Since we are using discrete samples, the boundary points will noticeably impact the integration results when the sample size is small. Therefore, half of the boundary points are incorporated in the integration and the other half are not.

Second, the integration over *J* is based on a uniform grid, which corresponds to the *x* values. However, the boundary points *x*_1_ and *x*_*n*_ (assume *x*_1_ and *x*_*n*_ are the largest and smallest value respectively) strongly deviate from this uniform assumption and the results will be affected when the sample size is small. We would like the integration to mimic the summation. Therefore, we compute the distribution of the largest sample (or smallest) and use the mean to get a new grid. The mean value *d* is computed by

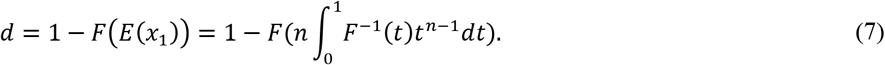

Here *t* should be densely sampled from 0 to 1. Then we get a new grid [*d*, …, *d* + (*i* − 1)(1 − 2*d*)/(*n* − 1), …,1 − *d*].

As mentioned above, in the presence of noise, the puncta from a certain threshold may contain holes (Fig. 2B). To make its shape more realistic, we may fill the holes (Fig. 2C). Besides, when we find the neighbors for each punctum, it is likely we will also include isolated pixels with higher intensity than the threshold (Fig. 2C). Our model is able to handle these effects since we allow some pixels in the candidate region to be lower than the threshold while some pixels in the background to be higher than the threshold. If we do not allow these exceptions and then the *M* pixels in the region is strictly brighter than all its *N* neighbors, our test statistic (Eq. 2) can be calculated by a truncated Gaussian model, which is less flexible in practice.

### 2.3 Iterative detection, FDR control, and post-processing

Our iterative detection and segmentation scheme is driven by the statistical significance of each region as computed above (Algorithm 1 and Fig. 1 D-I). Assuming the image is stored in 8 bits, we threshold the image with all intensity values (0 to 255). For each threshold *thr* ∈ {0,1, …,255}, we binarize the image *I* and get all connected regions in foreground. Suppose we totally get *K* regions with these 256 thresholds, the set of all such regions is denoted by *V* = {*S*_1_, …, *S*_*K*_}. We simply denote *S*_*k*_ as *k*, then *V* = {1, …, *K*}. We will build a tree *T*, whose nodes are *V*. We use *E* to denote the edge set describing the way to connect nodes (regions) in *V*.

Now each node *k* is associated with the region *S*_*k*_, along with the threshold *t*_*k*_ under which it is generated. Then the directed edge set is defined as *E* ≔ {(*i, j*)|*S*_*j*_ ⊆ *S*_*i*_, *t*_*j*_ = *t*_*i*_ + 1}, which means we link region *i* to a region *j* that is completely within it. However, not all inside regions should be linked. We link inside regions whose associated (more stringent) threshold is *t*_*i*_ + 1 (Fig. 1E). This structure is similar to conjunctive Bayesian networks (Beerenwinkel et al., 2007) and shares the similar principle as Mattes et al. (1999).

Each node *k* is also related to a neighbor pixel set *P*_*k*_ and a score *Z*_*k*_ from order statistics. Since the computation of order statistics depends on the choice of neighbor pixels, *Z*_*k*_ depends on *P*_*k*_. Recalling Eq.6, we have 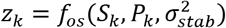. On one hand, *P*_*k*_ should include neighbor pixels of *S*_*k*_ and thus will be within an ancestry node of *k*, which is defined by the tree and denoted as *An*(*k*). The number of pixels in *P*_*k*_ needs to be carefully specified. If *P*_*k*_ is too large, many pixels far away from the candidate region *S*_*k*_ will be included and thus the comparison is not restricted to the local area. If *P*_*k*_ is too small, we lose the statistical power to assess the significance of the candidate region. We find that requiring *P*_*k*_ to have a similar size as the candidate region *S*_*k*_ is a good balance. In practice, we specify the neighbor region *P*_*k*_ by growing the candidate region *S*_*k*_ layer by layer until *P*_*k*_ is larger than *S*_*k*_. On the other hand, not all neighbor pixels of *S*_*k*_ should be included in *P*_*k*_ even though these pixels are close to *S*_*k*_, because these pixels may belong to another synaptic punctum region. Yet, a true synaptic punctum region should have a significant score. Therefore, we require *P*_*k*_ should not include any pixel of a significant region. Hence, *P*_*k*_ also depends on *z*_*k*_ as the significance of regions is determined by *z*_*k*_, which leads to the iterative scheme as described below.

**Algorithm 1.**
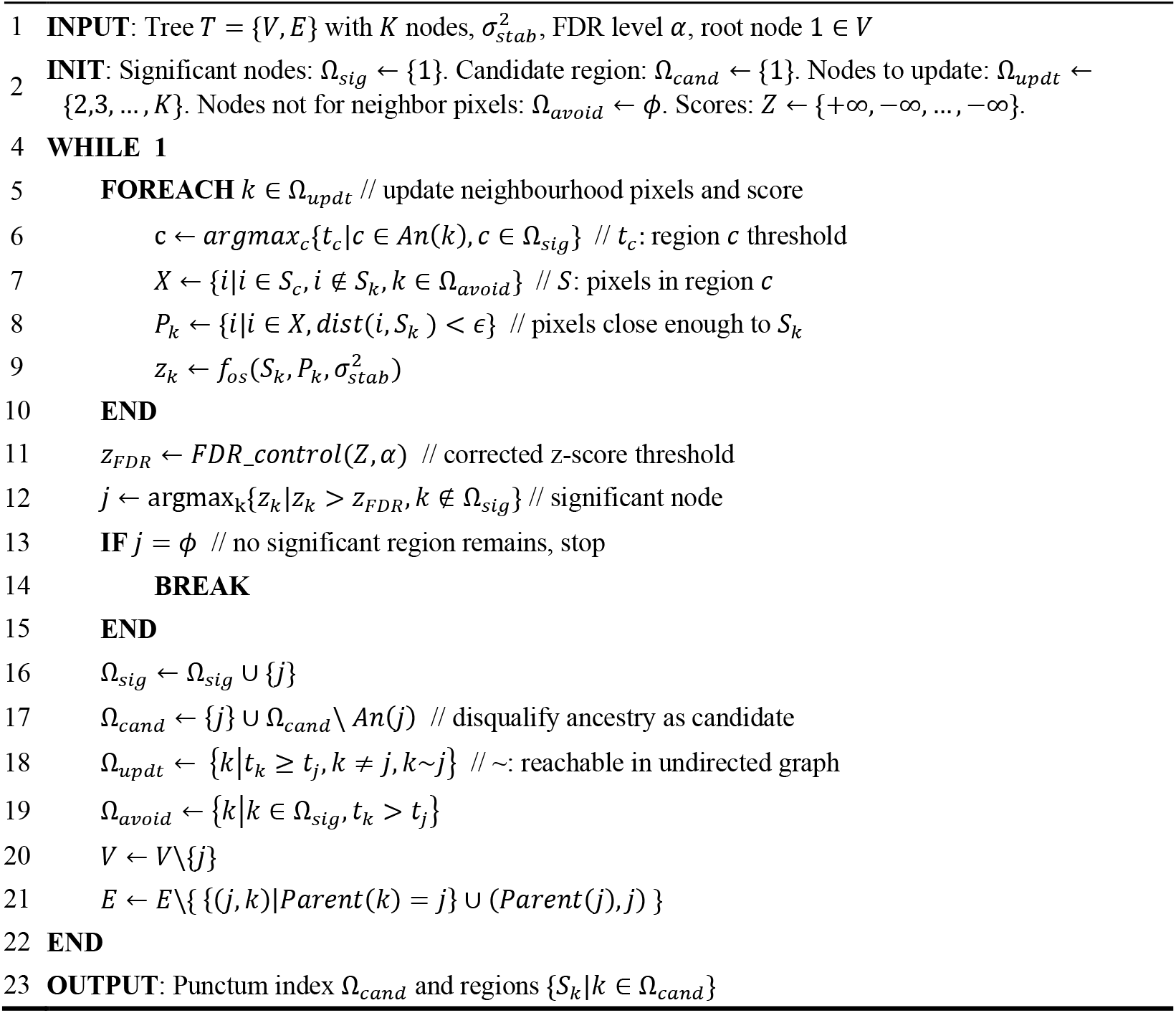
Order statistics based iterative detection and segmentation

Our algorithm iteratively updates *P*_*k*_ and *z*_*k*_ for each node *n* on tree *T*. We initialize the root node (*k* = 1, whole image) as the candidate region. For all other nodes, we initialize *z*_*k*_ = 0. All the other nodes now choose neighbor pixels *P*_*k*_ within the image (Fig. 1E) and do not need to avoid any pixels, because there is no significant region. Based on the choice of *P*_*k*_, we update *z*_*k*_ for all nodes (except for the root). Then we search for the most significant node *k* and update *P*_*k*_ for all the descendants of *An*(*k*), except those that are already significant (Fig. 1F). After that, node *k* is removed from the tree as a candidate synaptic punctum and its children will become the new root of a new tree (Fig. 1G). Again, the updated *P*_*k*_ will give us new *z*_*k*_. In later iterations, once any descendants of *k* becomes a new candidate, *k* is disqualified as a synaptic punctum. This drives the algorithm to avoid neurite-like structures (Fig. 1H-I).

With the mean and variance of order statistics under null hypotheses given in Section 2.2, we can calculate the scores of all relevant regions (Ω_updt_). We pick the one with the highest score and we need to determine whether we can add it to the list of significant regions (Ω_sig_). We want to keep the false discovery rate (FDR) lower than a given threshold among all synaptic puncta detected. The threshold (FDR level α) is a parameter specified by the user. A typical value is 0.05, which makes the false discovery rate lower than or equal to 5%. In each iteration, we learn from the FDR control procedure whether adding the newly selected region to the list of existing significant regions (Ω_sig_) will make the FDR lower than the threshold. Because overlapped regions may be correlated, we use the general case introduced by Benjamini and Yosef, (1995). Details are given in Supplementary Section 12. The total number of iterations depends on the number of synaptic puncta (significant regions) in the image and the user-specified FDR threshold. The algorithm stops when no more candidate region is determined as statistically significant. During the algorithm, no new region will be generated. The region marked as significant will not be updated later. Therefore, the maximum possible number of iterations is the number of regions from all thresholds and the algorithm is guaranteed to terminate.

Three rules based on the prior knowledge of the size and relative positions of puncta are applied to post-process the synaptic punctum candidates found by the algorithm (Uijlings et al., 2013). First, we filter out candidatesthat are too small or too large. Second, we expect the puncta to be close to circles or ellipses and we enforce this by setting threshold on the aspect ratio of puncta. Third, we expect the detected puncta to be roughly convex shaped, so we compare the area of the bounding box of a punctum with its area and remove those with low fill rate. In the experiments, these rules are applied to all methods. The remaining candidates are reported as synaptic puncta. The features of those detected puncta, such as size, brightness, and position, are then extracted.

The iterative detection algorithm is implemented on a component tree structure (Najman et al., 2006), which makes it possible to create the tree in quasilinear time instead of iterating over all threshold levels. When nodes are merged during the creation of the tree, features for test statistics of each node are recalculated the same way as in Ranefall et al. (2016).

### 2.4 Neurite quantification and statistical analysis

We use a steerable filter (Meijering et al, 2004) to trace the neurites. Unlike SynD (Schmitz et al., 2011), we relax the restriction that neurites must proceed from the soma to reduce false negatives for images with a large field of view. Details are given in Supplementary Section 13. This relaxation is desirable because such an image contains many neurons and some somas are not necessarily captured in it. Each detected neurite is cut into pieces. The intensity inside one piece is relatively homogeneous by cutting at branch points. Features including position, length, scale, and mean intensity for those pieces are extracted.

Each neurite piece has a set of synaptic puncta located on it or near it. In addition, as we discussed before, quite a few features of those puncta can be extracted. Therefore, we can build a model which uses each neurite piece along with those puncta on or near it as an input sample. This is a local feature analysis which is more informative than aggregating the feature across images. Also, for each image, we know the phenotype related to it, such as disease or normal conditions. Putting all these together, we can analyze the image with local synaptic puncta, neurites, and phenotypic features and discern the underlying associations.

### 2.5 3D implementation and support for pre- and post- synaptic channels

The idea of SynQuant can be naturally generalized to 3D images. The only difference is to define candidate punctum regions on 3D images after thresholding. We use the same way as used on 2D images, by selecting connected components. On 2D images, we use 4 or 8 connectivity to define connected components, while on 3D images, we use 10 connectivity, where we search 8 neighbors in XY plane and 2 directed neighbors in z direction. This is because for 3D images, the resolution of z-direction is usually lower than x- or y-direction.

We use a similar approach as PFSD (Simhal et.al., 2017) to combine the results from pre- and post-synaptic channels. For each punctum detected in the post-synaptic channel, we check all pre-synaptic puncta that overlap with it. Among them, we choose the one with the highest score and add it to the score of the post-synaptic punctum. We do not change the score of the post-synaptic punctum if no overlapping pre-synaptic puncta are found. This strategy is motivated by the fact the post-synaptic puncta usually have better co-localization with the gold standard synaptic cleft annotated in the electron microscopy channel (Collman et.al., 2015).

## 3 Results

We tested SynQuant on one simulation and three real data sets and compared it with four unsupervised methods and up to eleven variants of the supervised methods. The three real data sets include cultured cell (Mizuno et.al., 2018), multi-channel array tomography on brain slices (Collman et.al., 2015), and *in vivo* data (Bass et.al., 2017). We summarize the properties of the data sets in Supplementary Table 1. Among the methods we tested, SynD, PFSD, Bouton and DoGNet were designed for synaptic punctum detection, MS-VST and MP-HD are spot detection tools, and U-Net is a deep learning model for semantic segmentation. DoGNet contains two shallow neural networks models and two deeper models. Each model uses either an isotropic kernel or an anisotropic kernel to match the shape of puncta. For U-Net, we use the model provided in Kulikov et al., 2019. Here we did not include the two methods: SynPAnal and BGM3D, though they are mentioned in Table 1 and Table 2. The reason for excluding SynPAnal is that it is a semi-supervised method which needs user to manually crop the region of dendrite first. BGM3D is a method based on global thresholding like SynD, so we chose one of them in our comparison. Further, experiments were performed to measure the influence of Down syndrome on synaptic density while taking all confounding factors (extracted features) into consideration (Supplementary Section 7). We implemented the algorithm in Java as a Fiji plugin (Schindelin, Johannes, et al., 2012). Our implementation supports both 2D and 3D data, as well as combining pre-synaptic and post-synaptic channels. All the data sets, labels, and code to generate the synthetic data are also available on the GitHub website.

We evaluated the performance by precision-recall curve. We use Intersection-over-Union (IoU) to infer true positive (TP), which is widely used in object detection problems. If the overlap of ground truth and the detected punctum is larger than 50% of their union, the detected punctum is viewed as a TP. IoU is more suitable when we want to jointly evaluate the detection and segmentation performance. For real data, we do not have pixel level annotations for puncta, so we set the threshold to 0%, that is, a TP is claimed as long as the detection has any overlap with the ground truth or annotation. Precision is defined as the TP/(TP+FP) and recall is defined as TP/(TP+FN), where FP is number of false positive and FN is false negative. The F score is 2 × precision × recall/(precision + recall).

Each method will provide a score map with the same size as the input data, we threshold the score map from its minimum value to its maximum value with 100 thresholds. We calculate a precision-recall pair for the puncta above each threshold. As SynD does not associate a score with the puncta, we simply use all detected puncta. As Bouton only reports the center of each punctum, we replace the center by a square with similar size as the average actual size of puncta in the data. DoGNet provides a score map as well. In the paper, authors hard threshold the score map, find local maximum and suppress nearby ones. To make fair comparison, we treat it the same way as all other methods. Actually, their hard thresholding based post-processing in DoGNet makes its performance worse in our experiments. We report the best F1 score for all points in the precision-recall curve. We also calculate the average precision based on the precision-recall curve (Everingham et al., 2010), which scans recall from 0.01 to 1, with step size 0.01. Average precision summarizes the information contained in the precision-recall curve and is a more comprehensive measure than the best F1 score.

Some methods require the choice of parameters and/or manual intervention. SynD requires manual selection of a global threshold. We use the ground truth to automatically determine a threshold. Specifically, the threshold should lead to the highest correlation with ground truth after binarization. The scales for MS-VST and the smoothing parameter for MP-HD are tuned for the best performance in F1 scores. The parameters of Bouton are changed to consider the puncta sizes. For DoGNet and U-Net, we use the same network architectures and hyper parameters as provided in Kulikov et.al., 2019. We show the results of four models proposed in Kulikov et.al., 2019, which include two shallow networks and two deep networks. Each network either uses isotropic filter or anisotropic filter.

To test supervised methods, we use part of the labeled data for training. We use ‘-trained’ suffix for each supervised method to indicate that it is trained from scratch using these data. The remaining labeled data are used to test all methods. For Bouton, we also use the pre-trained model, which we denote as ‘Bouton-pretrained’. For DoGNet, we also try to first train the Collman’s data, which has the largest number of labels, and then fine tune it using the labeled data provided in each data set. This allows us to train the deeper versions of models in DoGNet. More details are provided in the sections for each data set. In all experiments, post processing is applied. The minimum size of a punctum is 8 voxels (except for the synthetic data, where the minimum size is 4); the maximum size is 300 voxels; the aspect ratio should be between 0.5 and 2; the ratio of voxels to bounding boxes should be larger than 0.5. These rules are applied to all methods tested.

### 3.1 Results on synthetic data

Our simulation data consists of both synapse and neurite like signals to mimic realistic data (Supplementary Section 3), which simulates the punctum inhomogeneity and antibody non-specificity. We compared the F1 score of all methods with different simulation settings. We first simulated the impact of Poisson Gaussian noise on the performance (Supplementary Section 4.1). The SNR was calculated as the average SNR for each simulated punctum. For all SNRs, SynQuant performs always the best. For example, when IoU threshold is 0, the best F1-score of SynQuant outperforms the best performing peer method by 0.162 (0.981 vs. 0.819) under 11.5 dB SNR. Then we studied the impact of the range of punctum size (Supplementary Section 4.2). SyQuant still performs the best for all experiments. When the range of punctum size is 9 to 150 pixels, the best F1-score from SynQuant outperforms the best peer method by 0.159 (0.962 vs. 0.803) when the IoU threshold is 0.5 and SNR is 17.2 dB. With IoU threshold equals to 0.5, inaccurate segmentation of the puncta will be considered as false positive. The experiments in the synthetic data show that SynQuant is robust to noises and punctum size changes. More details can be found in the Supplementary Section 4.

### 3.2 Results on Bass’ 3D *in vivo* data

We tested SynQuant on the *in vivo* 3D image data available in Bass et.al., 2017. In this data, signals can be observed on both neurites and synaptic puncta (Fig 1A). The data set contains 100 3D image sets. 80 of them are partially labeled for training Bouton model. The remaining 20 are completely annotated for testing. As the training for DoGNet and U-Net need all puncta to be labeled, we cannot use the 80 training images. Instead, we divide the 20 well annotated images into two groups. The first 10 are used for training DoGNet and U-Net and the remaining 10 are used for the evaluation of all methods. Each image set has size 512 by 512 in XY plane. The number of Z stacks varies from 17 to 50. The resolution on X-Y direction is 0.147 *μm* and 1 *μm* for Z-direction. Though the data is 3D, the annotations are 2D bounding boxes of the puncta. Besides, we find these labels are all oversized (see Supplementary Fig.10), which contain many redundant background pixels. Nearby puncta are easily overlapped with each other with these large labels. However, by examining the data, one punctum always occupies single isolated spatial location. Thus, we reduce the label size by taking the center of each punctum and put a square whose size is similar to the average actual size of puncta in the image.

For Bouton, we used the pre-trained model based on the 80 training images and did not need to train it by ourselves. In Bouton, the images are mean projected to 2D before analysis. For SynQuant and PFSD, we directly detected in 3D. For MS-VST and MP-HD, the performance of 3D version is comparable with 2D version, so we only show the 2D results here. As we do not have 3D labeling for this data set, we use DoGNet in mean projected image as well. The 10 training images are not sufficient to train the deeper models in DoGNet and cannot make correct predictions (N/A in Table 3). Therefore, other than directly training deeper models, we also used the 10 images to fine tune the deeper DoGNet/U-Net models that are pre-trained in Collman’s data.

**Table 3.**
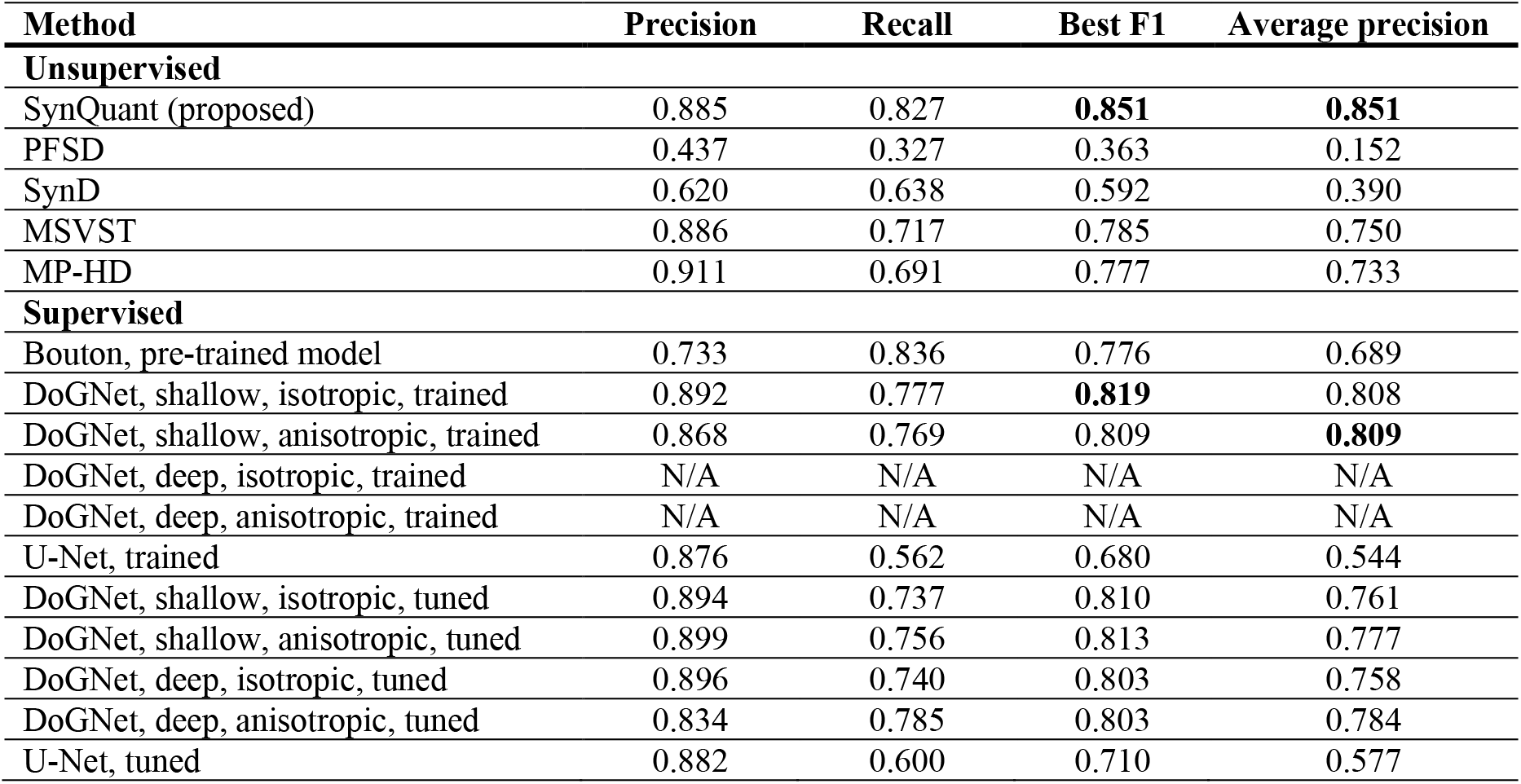
Results on Bass’ in vivo 3D data

Results show that for both F1 score and average precision, SynQuant performs the best among all methods compared. DoGNet is the next best performing method. Bouton fails to detect the center of puncta accurately, which degrades its performance.

### 3.3 Results on Collman’s array tomography data

We tested SynQuant on the array tomography data provided in Collman et.al., 2015. There are two data sets provided: Collman 14 data with 39 z-stacks and Collman 15 data with 27 z-stacks. Each data is stained with multiple antibodies. We use the PSD-95 stained post synaptic channel and the Synapsin I labeled pre-synaptic channel. Each data is also imaged with electron microscopy (EM). The synaptic clefts in the EM images were annotated. The annotations are down-sampled to match the original resolution of the fluorescence staining (0.1 *μm*/pixel). We use Collman 14 data to train all supervised methods and test on Collman 15 data. The ground truth annotation is in EM channel, some of which do not correspond to the puncta in synaptic channels. This kind of inconsistency usually happens when the imaging field of view for fluorescence channels and EM channel are different. To correct it, we check each annotation, if it does not have any fluorescence staining co-localized, we remove that annotation.

The annotations on EM channel are not suitable for training the model in Bouton, so ‘Bouton-trained’ cannot make correct prediction if trained in this way. SynD does not support 3D data, so we do not list it here, while SynQuant and PFSD can be directly applied on 3D data. For other methods, we detect puncta stack by stack and combine the score maps afterwards. This is the default approach used by DoGNet and was shown to perform better than 3D version of DoGNet (Kulikov et.al., 2019). The 3D version of MS-VST and MP-HD also performs worse than their 2D version. Since the Collman 14 data has the largest number of ground truth labels, we do not use other data to train first. Therefore, we do not have the ‘tuned’ models listed in Table 4. While DoGNet and PFSD are able to integrate information from two channels, other peer methods do not have this functionality. Therefore, for these methods, we apply the same method SynQuant uses to combine results from the pre-synaptic and the post-synaptic channels. We evaluate the performance on pre-synaptic channel, post-synaptic channel, and combined results. SynQuant performs best among all unsupervised methods and the results are comparable with the best performing supervised methods. In terms of average precision, SynQuant performs the best among all methods. Detailed results on pre-synaptic and post-synaptic channels can be found in the Supplementary Section 5. Our method is developed to detect synaptic puncta, which do not have a one-to-one correspondence with synaptic cleft, but achieves comparable performance with DoGNet.

**Table 4.** Results on Collman’s array tomography data

| Method | Precision | Recall | Best F1 | Average precision |
| --- | --- | --- | --- | --- |
| **Unsupervised** |  |  |  |  |
| SynQuant (proposed) | 0.882 | 0.699 | **0.780** | **0.754** |
| PFSD | 0.885 | 0.589 | 0.707 | 0.666 |
| MSVST | 0.905 | 0.648 | 0.756 | 0.715 |
| MP-HD | 0.876 | 0.648 | 0.745 | 0.680 |
| **Supervised** |  |  |  |  |
| Bouton, pre-trained model | 0.602 | 0.224 | 0.327 | 0.229 |
| Bouton, trained | N/A | N/A | N/A | N/A |
| DoGNet, shallow, isotropic, trained | 0.795 | 0.691 | 0.739 | 0.638 |
| DoGNet, shallow, anisotropic, trained | 0.868 | 0.636 | 0.734 | 0.621 |
| DoGNet, deep, isotropic, trained | 0.880 | 0.708 | **0.784** | **0.704** |
| DoGNet, deep, anisotropic, trained | 0.897 | 0.665 | 0.764 | 0.691 |
| U-Net, trained | 0.823 | 0.631 | 0.715 | 0.626 |

### 3.4 Results on neuron-astrocyte co-cultured data

We tested SynQuant and other methods on our in-house neuron-astrocyte co-culture data set, which contains 16 images. The size of each image is 256 by 256 pixels. Each image contains two channels: the synapse channel (green) labeled by Synapsin I and the neurite channel (red) labeled with Tuj1. We use the synapse channel for the comparison of synaptic punctum detection and manually labeled the synapse channels in these 16 images. Only the puncta that are clear enough to reach the consensus among two experts are considered as ground truth. 8 images are used for training supervised methods and the remaining 8 are used for testing all methods.

For Bouton, we use both the pre-trained model from Bass’ data and the model trained from scratch on these 8 training images. For DoGNet and U-Net, directly using the model pre-trained on Collman’s data does not perform well, so we fine tune the model pre-trained on Collman’s data using these 8 training images. We also directly train the DoGNet and U-Net models from scratch using the 8 training images. The deeper models in DoGNet cannot be successfully trained given limited training data, though they can be trained first with the larger Collman’s data and tuned after that. Even though U-Net is not designed for synaptic punctum detection, it also works well in this data.

We report the performance based on the average of the 8 test images. The precision and recall in Table 5 correspond to the best F1 score. SynQuant outperforms all unsupervised methods and is comparable with the best supervised method in terms of F1 score. In terms of average precision, SynQuant outperforms all the other methods.

**Table 5.** Results on neuron-astrocyte co-cultured data

| Method | Precision | Recall | Best F1 | Average precision |
| --- | --- | --- | --- | --- |
| **Unsupervised** |  |  |  |  |
| SynQuant (proposed) | 0.942 | 0.878 | **0.904** | **0.933** |
| PFSD | 0.464 | 0.678 | 0.536 | 0.336 |
| SynD | 0.299 | 0.941 | 0.434 | 0.276 |
| MSVST | 0.919 | 0.803 | 0.853 | 0.861 |
| MP-HD | 0.925 | 0.765 | 0.835 | 0.822 |
| **Supervised** |  |  |  |  |
| Bouton, pre-trained model | 0.550 | 0.549 | 0.533 | 0.451 |
| Bouton, trained | 0.567 | 0.519 | 0.494 | 0.430 |
| DoGNet, shallow, isotropic, trained | 0.917 | 0.866 | 0.888 | 0.880 |
| DoGNet, shallow, anisotropic, trained | 0.934 | 0.812 | 0.882 | 0.883 |
| DoGNet, deep, isotropic, trained | N/A | N/A | N/A | N/A |
| DoGNet, deep, anisotropic, trained | N/A | N/A | N/A | N/A |
| U-Net, trained | 0.931 | 0.861 | 0.892 | 0.889 |
| DoGNet, shallow, isotropic, tuned | 0.929 | 0.883 | **0.903** | **0.894** |
| DoGNet, shallow, anisotropic, tuned | 0.912 | 0.868 | 0.889 | 0.885 |
| DoGNet, deep, isotropic, tuned | 0.932 | 0.846 | 0.882 | 0.859 |
| DoGNet, deep, anisotropic, tuned | 0.908 | 0.759 | 0.824 | 0.772 |
| U-Net, tuned | 0.828 | 0.847 | 0.882 | 0.873 |

We also applied SynQuant to study the relationship between synaptic punctum density on neurites and Down-syndrome cell line using this in-house data set. Details can be found in Supplementary Section 7.

### 3.5 Summary of the experimental results

In summary, tested on a large variety of experiment settings, including 2D vs 3D, single vs multiple channels, confocal, two-photon or array tomography, neurite contamination vs no contamination, and manual labeling vs EM annotation, SynQuant always outperforms other state-of-the-arts in terms of the average precision, which is a preferred measure by researchers over the best F1 score. This is because the best F1 score is sensitive to noise and cannot indicate the robustness of algorithms. However, even using the best F1 score, SynQuant still performs the best among all unsupervised methods and achieves better or comparable performance compared with the supervised methods. It is worth mentioning that none of the supervised methods can compete with SynQuant consistently across all these data sets, that is, their performance varies across different data sets. For instance, the DoGNet model (shallow, isotropic, tuned) performes comparably to SynQuant in the co-culture data in terms of the best F1 score (SynQuant 0.904 vs DoGNet 0.903), however, the same model is much worse than SynQuant in Bass’ data (SynQuant 0.851 vs DoGNet 0.810) and in Collman’s data (SynQuant 0.780 vs DoGNet 0.734). Furthermore, the ground truth of the real data we experimented on can only provide location information of puncta without pixel-wise labels. Thus, we cannot measure and compare the pixel-level segmentation accuracy, which is a strong point of SynQuant as we can see from the results of synthetic data set.

## 4 Discussion

We have presented a new automatic synapse quantification framework (SynQuant) for segmentation and quantification of heterogeneous and noisy images of synapses and dendrites. SynQuant is able to detect and segment synaptic puncta accurately. It can extract comprehensive features from both synaptic puncta and neurites and analyze their relationships.

The superior performance of SynQuant comes from the effective utilization of the local region-neighbor information. Enjoying the same principle as Hariharan et al. (2014), the probability principled iterative detection and segmentation algorithm uses the tree structure of regions to choose the correct neighborhood pixels. Order statistics provide an unbiased score to indicate the likelihood that one candidate region is a true synaptic punctum. The choice of neighborhood pixels and the computation of order statistics are iterated until no statistically significant regions can be found in the tree structure. Compared with existing spot detection methods, SynQuant is able to extract accurate segmentation results, which allows access to important features for synapse studies. The main parameter needed for SynQuant is the FDR level to be controlled and the algorithm reports the p-value or z-score for each region detected.

Incorporated with properties extracted from the neurite channel, effects of cell types and disease phenotypes on synaptic punctum density can be revealed. With a large field of view, the number of neurons involved is large. As a result, different neurons and their corresponding neurites and synapses are at different development stages due to cell type and other stochastic effects. As such, these neurites and synapses must be treated locally and we should not aggregate the features in the image. In SynQuant we use a segmented neurite piece as a sample in the regression analysis. Thus, the relationships we found are more comprehensive by including these local confounding factors and can more reliably reveal the relationships between disease and synaptic features.

SynQuant makes it easier to use antibodies that are not so specific without the requirement of creating training labels. Though supervised methods (especially DoGNet) work well on the data set we tested above, there are several things to emphasize. First, the creation of training labels can be time consuming, especially if a lot of training samples are needed for better performance. Second, the model trained based on existing labels usually cannot be directly applied to another data, unless the data sets are obtained under very similar experiment setups. Therefore, more training labels on the new data set are needed. Third, supervised models are unavoidably influenced by human bias introduced in data labels. For example, we observed that in the three real data sets, synaptic puncta with low intensities are much easier to be missed in the ground truth than those brighter ones. Under such biased labels, supervised models have a high risk of missing dimmer puncta.

SynQuant supports 3D data as well as multi-channel data with both the pre-synaptic puncta and post-synaptic puncta. Moreover, SynQuant is a general framework to analyze images with a high level of non-specificity. We can naturally adapt and apply it to bio-medical images beyond synapse staining, such as particle detection for the particle tracking problem.

## Supplementary material

### 1 Overall framework of SynQuant

As shown in Supplemental Figure 1, SynQuant is designed to detect and quantify synaptic puncta and related features. It can also trace neurites and extract features from them. If some images are acquired when cells are under normal conditions and some images come from cells under disease conditions, the features extracted by SynQuant on each image can be used for subsequent statistical analysis to reveal the impact of synapse on the disease.

**Supplemental Figure 1.**
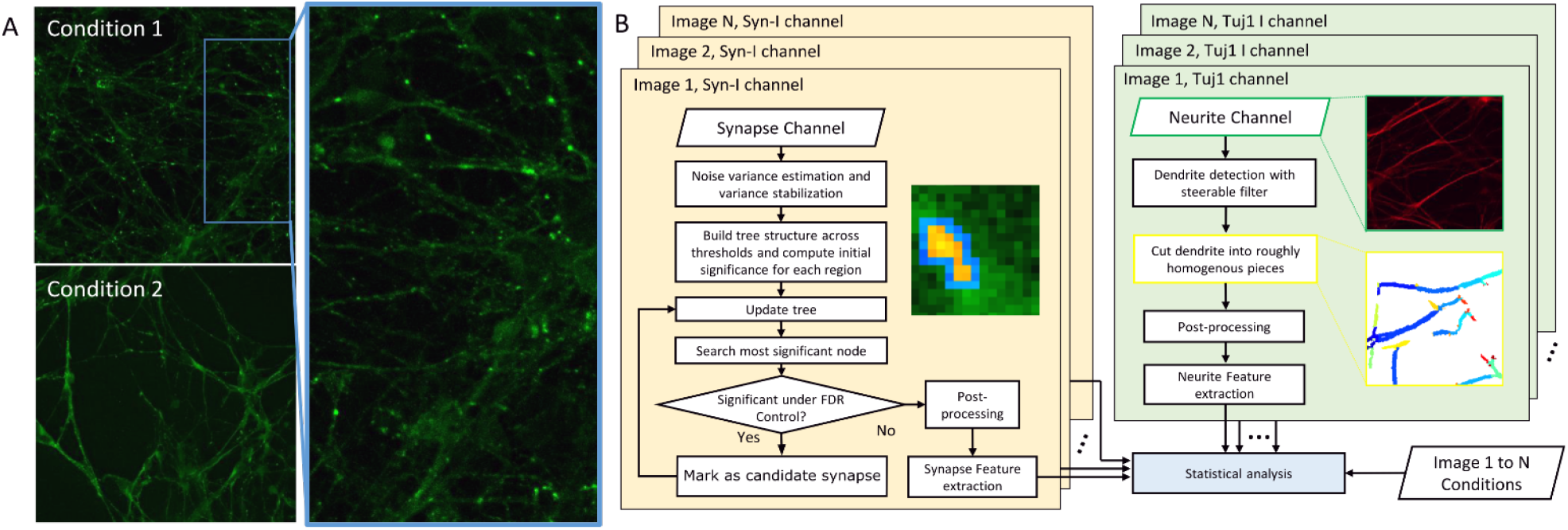
(A) Synapsin I labeled channel acquired by confocal microscopy. Green dots are potential synaptic puncta. Two images on the left panel belong to control (top) and case (bottom) groups, respectively. (B) Flowchart of SynQuant. Left panel: the synapse detection algorithm. The inset figure shows the choice of neighborhood pixels (blue) for a region (yellow pixels). Right panel: neurite features. Top sub figure shows the Tuj1 stained neurite channel. The bottom one shows the local homogenous cuts on the traced neurites. The extracted puncta and neurite features from N images are used for statistical analysis between conditions.

### 2 Summary of data sets

The descriptions of the data sets used in the experiments are shown in Supplemental Table 1. There are 100 images in Bass’s data (Bass et.al., 2017), but only the 20 test images are well labeled. The synthetic data are generated based on the dendrite information obtained from the neuron-astrocyte co-cultured data (Mizuno et.al., 2018). In Collman’s data (Collman et.al., 2015) downloaded from https://neurodata.io/, the fluorescence staining channel was up-sampled to match the resolution of the electron microscopy channel. Here we down-sample them to their original 100 nm XY spatial resolution.

**Supplemental Table 1.** Summary of data sets used in the experiments

| Name | Width and Height (pixels) | Z stack number | nm/pixel on XY | Gap on Z (nm) | Channels | Technique | Image number |
| --- | --- | --- | --- | --- | --- | --- | --- |
| **Bass**<br>(Bass et.al., 2017) | 512, 512 | 17 to 50 | 147 | 1000 | GFP | In vivo two-photon | 100 |
| **Co-culture**<br>(Mizuno et.al., 2018) | 1024, 1024 | 1 | 208 | N/A | Synapsin I | Confocal | 40 |
| **Collman 14**<br>(Collman et.al., 2015) | 368, 295 | 39 | 100 | 70 | Multiple | Array tomography | 1 |
| **Collman 15**<br>(Collman et.al., 2015) | 141, 101 | 27 | 100 | 70 | Multiple | Array tomography | 1 |
| **Synthetic** | 1024, 1024 | 1 | 208 | N/A | N/A | N/A | 90 |

### 3 Synthetic data generation

We generate synthetic data where the puncta have different brightness and size. The anti-body is also not perfectly specific, which also shows signals in the neurite like structures. We design simulation to have different signal to noise ratio (SNR). We also change the puncta size range since it is commonly observed in Bass’s 3D in vivo data (Bass et.al., 2017) and our in-house neuron-astrocyte co-cultured data that the puncta size varies significantly. Therefore, a good algorithm should also be able to handle different noise levels and various puncta sizes.

We simulated neurite-like signals by putting the segments of traced branches into the blank simulation image. These branches are obtained from the Tuj1 labeled channel in the in-house neuron-astrocyte co-cultured data. After that, the image was smoothed by Gaussian filter. Then synapses are randomly cast on the simulated dendrite with various shapes. The synapse intensity is assumed to be greater than the corresponding dendrite. Finally, Poisson-Gaussian noise was added.

More specifically, the simulation data is generated with four steps. 1) To fully simulate the conditions of neurite, we first extract the neurite from a neuron-astrocyte co-cultured data in control group (neurite extraction method is discussed in Supplemental Text Section 13). Based on the results of extraction, we cut them into small pieces. Pixels inside each piece are viewed roughly homogeneous. 2) Generate neurite pieces on our simulation image at the same positions as the real data, with scales and brightness also similar to the ones on the real image. For each neurite piece, we use a Gaussian kernel with standard deviation of 1 to smooth it. 3) Cast synapses on each neurite piece. The number of synapses is based on the synapse numbers obtained from real data on that branch. The synapse intensity is decided by the summation of three values: intensity of the neurite it falls on, a contrast parameter and a random noise value. The contrast parameter is used to define average contrast across the image between synapses and neurites they fall on. The random value is generated by multiplying the contrast parameter with a standard Gaussian random variable. The minimum size of the puncta is set to 9 while the maximum sizes of the puncta are changed in different simulation setups. In one image, the size of the puncta is randomly chosen between the minimum size and the maximum size. 4) Mixed-Poisson-Gaussian noise is added to the image. The noise is added using the ‘imnoise’ function of MATLAB with the ‘localvar’ option. The noise level corresponding to the minimum intensity is not changed and the noise level for the maximum intensity is changed to generate different signal to noise ratios. After adding noise, we calculate the signal to noise ratio for each simulated punctum and use their average as the SNR for the whole image.

We generate synthetic images with three different SNRs and three different ranges of puncta size, so there are nine simulation setups. In each setup, we repeated the simulation for 10 times. The total number of images simulated is 90. Examples of synthetic data can be found in Supplemental Figure 2 and Supplemental Figure 3.

**Supplemental Figure 2.**
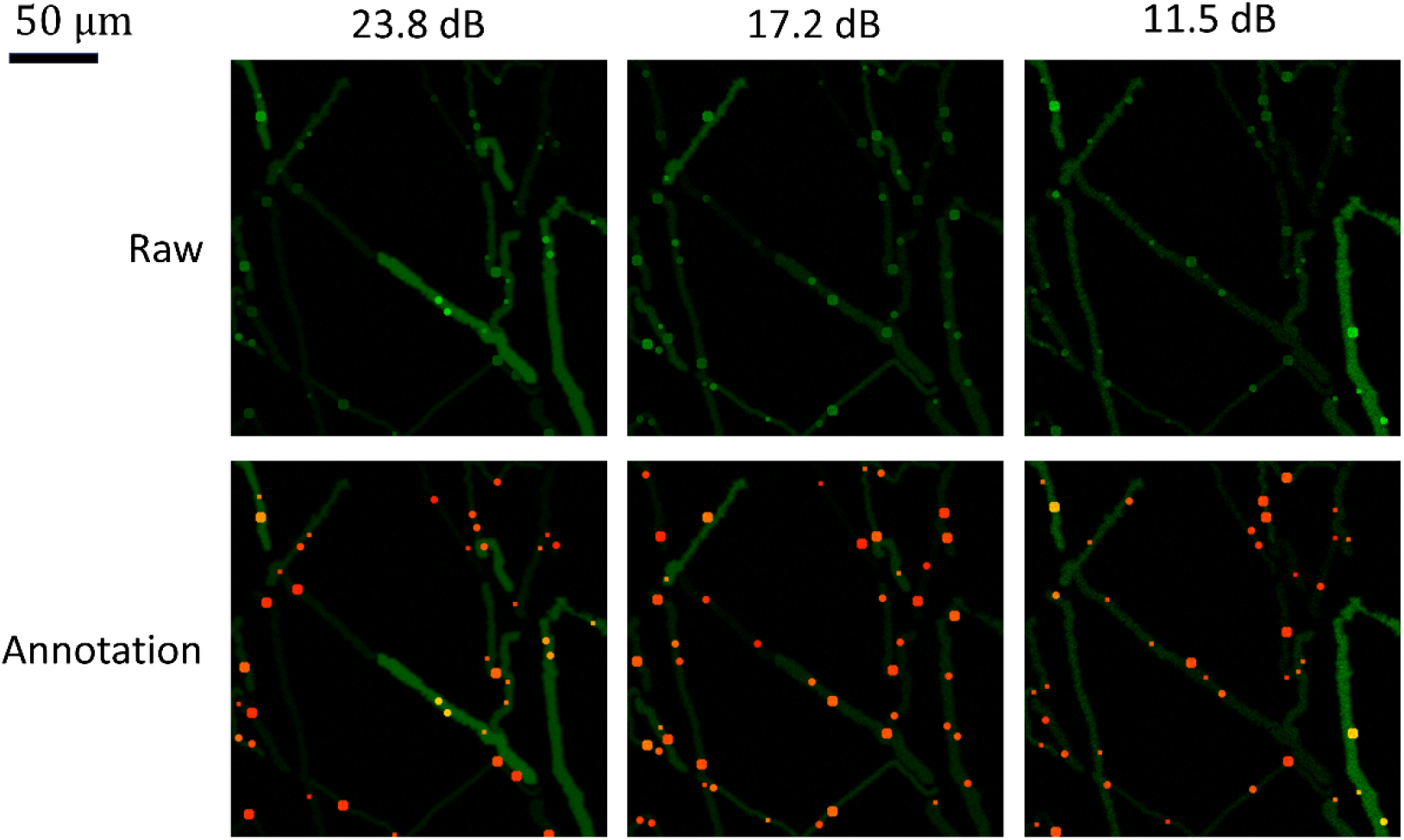
Synthetic data example at different signal to noise ratios. The puncta size ranges from 9 pixels to 50 pixels. The top row is the raw data. The bottom row shows the raw data overlaid with ground truth labels, which are shown in red. The size of the original image is 1024 by 1024 pixels. Here a patch of 256 by 256 pixels is shown for each image.

**Supplemental Figure 3.**
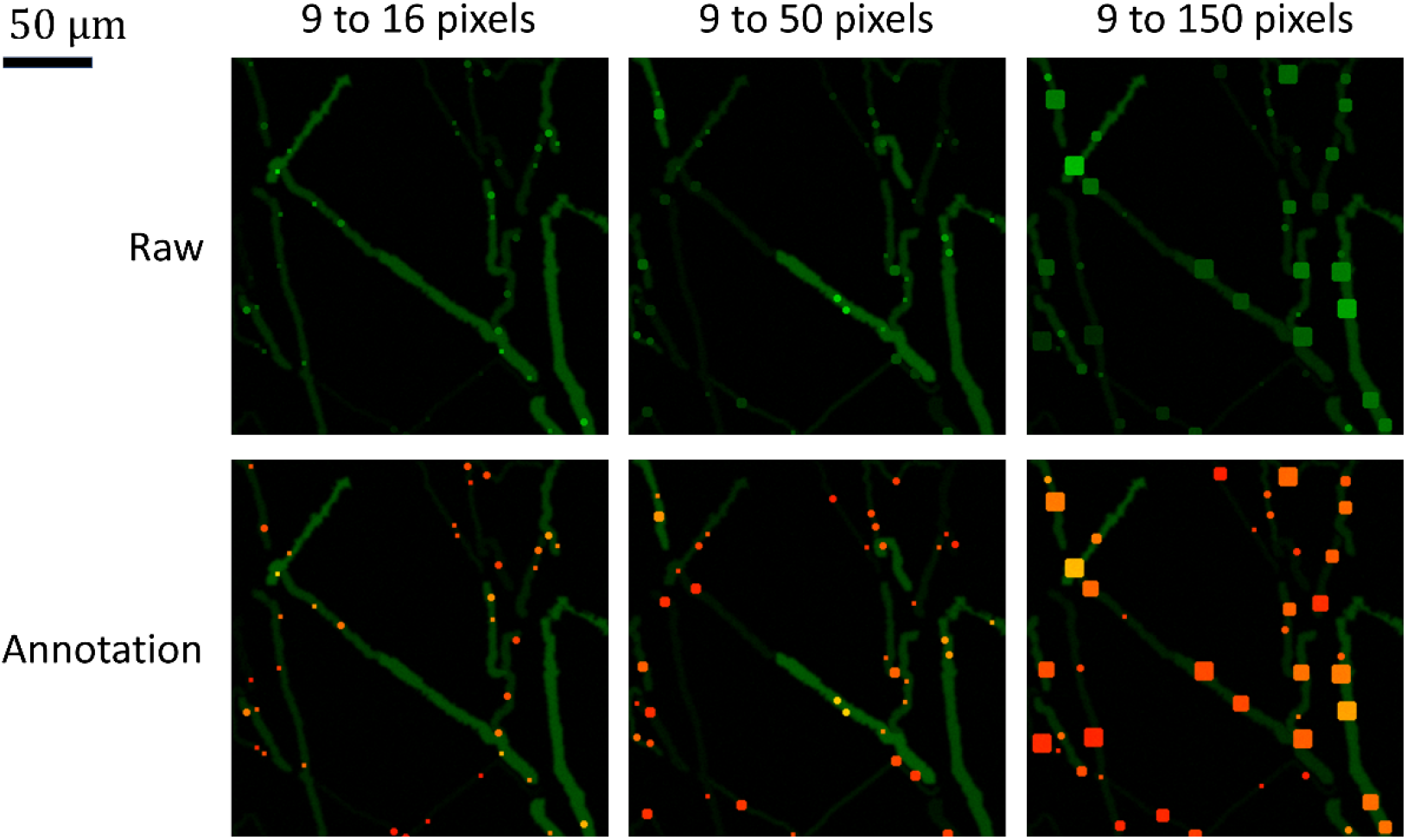
Synthetic data example at different ranges of puncta sizes. The signal to noise ratio is 23.8 dB. The top row is the raw data. The bottom row shows the raw data overlaid with ground truth labels, which are shown in red. The size of the original image is 1024 by 1024 pixels. Here a patch of 256 by 256 pixels is shown for each image.

### 4 Results on synthetic data

We studied the impact of noise levels and the range of puncta size on the detection accuracy. We also inspected the performance of SynQuant under different thresholds of the intersection of union (IoU). As we commonly observed in Bass’s 3D *in vivo* data (Bass et.al., 2017) and our in-house neuron-astrocyte co-cultured data, the size of puncta can change greatly in the same image. In addition, the noise level of data obtained from different technologies and different experiments differs greatly and the detector should be able to work well under different noise levels and puncta size ranges.

As in the experiments with real data, we used a low IoU threshold to study the performance of puncta detection. In simulation data, we also used a higher IoU threshold to simultaneously study the performance of detection and segmentation, which was not feasible in real data due to the lack of pixel level labeling in the fluorescence staining.

More specifically, an IoU threshold of 0 means that, as long as a detected punctum overlaps with a ground truth punctum, it will be considered as detected. We did not duplicate counting. For example, if a ground truth punctum was found to be overlapped with one punctum detected, it would be marked as used and other detected puncta could no longer overlap with it. This threshold mainly measures the quality of detecting puncta. If the IoU threshold is 0.5, only when the ratio of the overlapping area of the detected puncta and the ground truth puncta to the union of their area is larger than 0.5, the puncta was considered as detected. This measures not only the detection accuracy, but also the quality of segmenting puncta. The precision-recall curve under one experiment setup with two different IoU thresholds is also given later in this section to illustrate the performance differences under these two IoU thresholds.

The anisotropic version of deep DoGNet could not be successfully trained in this simulation. As Bouton does not segment synapse, we did not compare with it in experiments where the IoU threshold is 0.5. It was still used when the IoU threshold is 0 where we mainly studied the performance of puncta detection. Each puncta center detected in Bouton was replaced by a square of 5 by 5 pixels before performance evaluation.

#### 4.1 Impact of noise level on detection accuracy

We show the results of experiment where the SNR changes (Supplemental Figure 4 and Supplemental Figure 5). The size of puncta ranges from 9 to 50 pixels. It is clear that SynQuant outperforms all peer methods in this simulation in all signal to noise ratios tested. The next best performing method is DoGNet, though its advantage over MS-VST is not too large. The relative performance of different models in DoGNet also changes in different experiment setups. With larger noise level, the detection performance of all methods degrades. The performance of most methods also drops when the IoU threshold is 0.5 while SynQuant suffers much less from this more stringent criterion.

#### 4.2 Impact of puncta size range on detection accuracy

We study the impact of puncta size range on the performance (Supplemental Figure 6 and Supplemental Figure 7). The SNR in this experiment was kept at 23.8 dB. When size range can only change from 9 pixels to 16 pixels, DoGNet, MS-VST, and MP-HD performs well, especially when the IoU threshold is 0. Performance of all peer methods drops significantly when the puncta are allowed to have more diverse size range, which is a common phenomenon in real data. In contrast, the performance of SynQuant is not significantly influenced by the increasing range of puncta sizes. When IoU threshold is 0, the performance of Bouton becomes better when puncta size is allowed to be within a larger range. This is mainly because Bouton cannot detect the center of puncta accurately. When a larger proportion of the puncta have larger size, it is more likely for the detected centers to be inside the ground truth puncta.

#### 4.3 An example precision recall curve

We show the precision-recall curves for all methods when applied to a synthetic data (Supplemental Figure 8 and Supplemental Figure 9). In that data, the puncta size changes from 9 pixels to 50 pixels and the SNR is 17.2 dB. Each method corresponds to a precision-recall curve. The different points in the curve are obtained by changing the thresholds of the score map estimated by that method. If the score map ranges from 0 to 1, we set 99 thresholds from 0.01 to 0.99. For each threshold, the connected components in the binarized score map are considered as detected puncta. SynD does not assign scores to the puncta it detected, so its precision-recall curve has only one point. For each method, the precision and recall that correspond to the best F1 score are shown as a dot with the same color as the curve for that method. When the IoU threshold is 0.5, Bouton has very low performance as it is not designed to segment puncta. A precision-recall curve is not necessarily to be monotonic since increasing the threshold in the score map may lead to results that reduce precision. In this simulation, SynQuant outperforms all other methods. DoGNet, U-Net, MS-VST, and MP-HD perform better than other peer methods. The performance advantage of SynQuant is more obvious when the IoU threshold is set to 0.5.

**Supplemental Figure 4.**
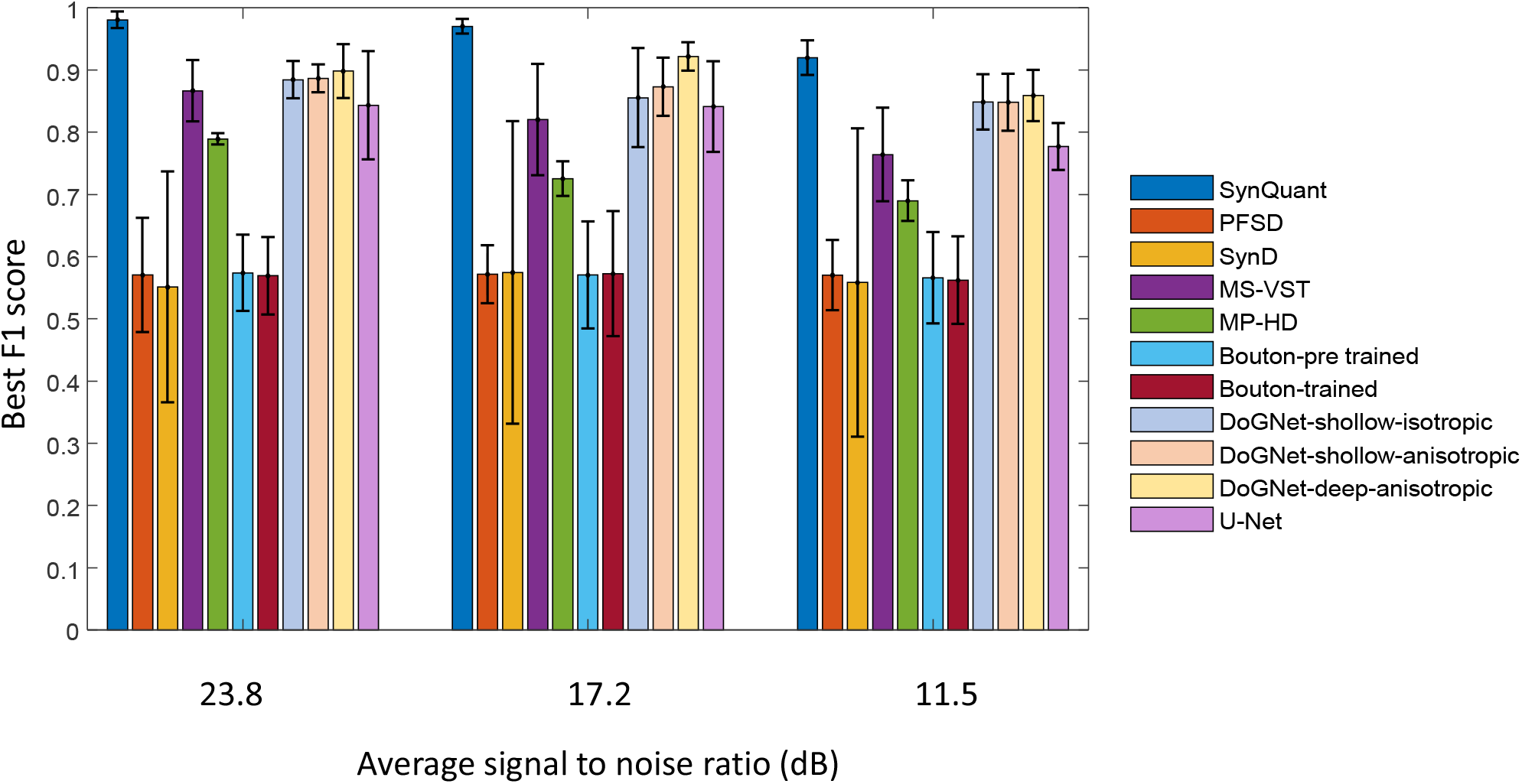
Performance on simulation data with different signal to noise ratios. The range of puncta size is from 9 to 50. The IoU threshold is 0. The error bars represent 95% confidence interval.

**Supplemental Figure 5.**
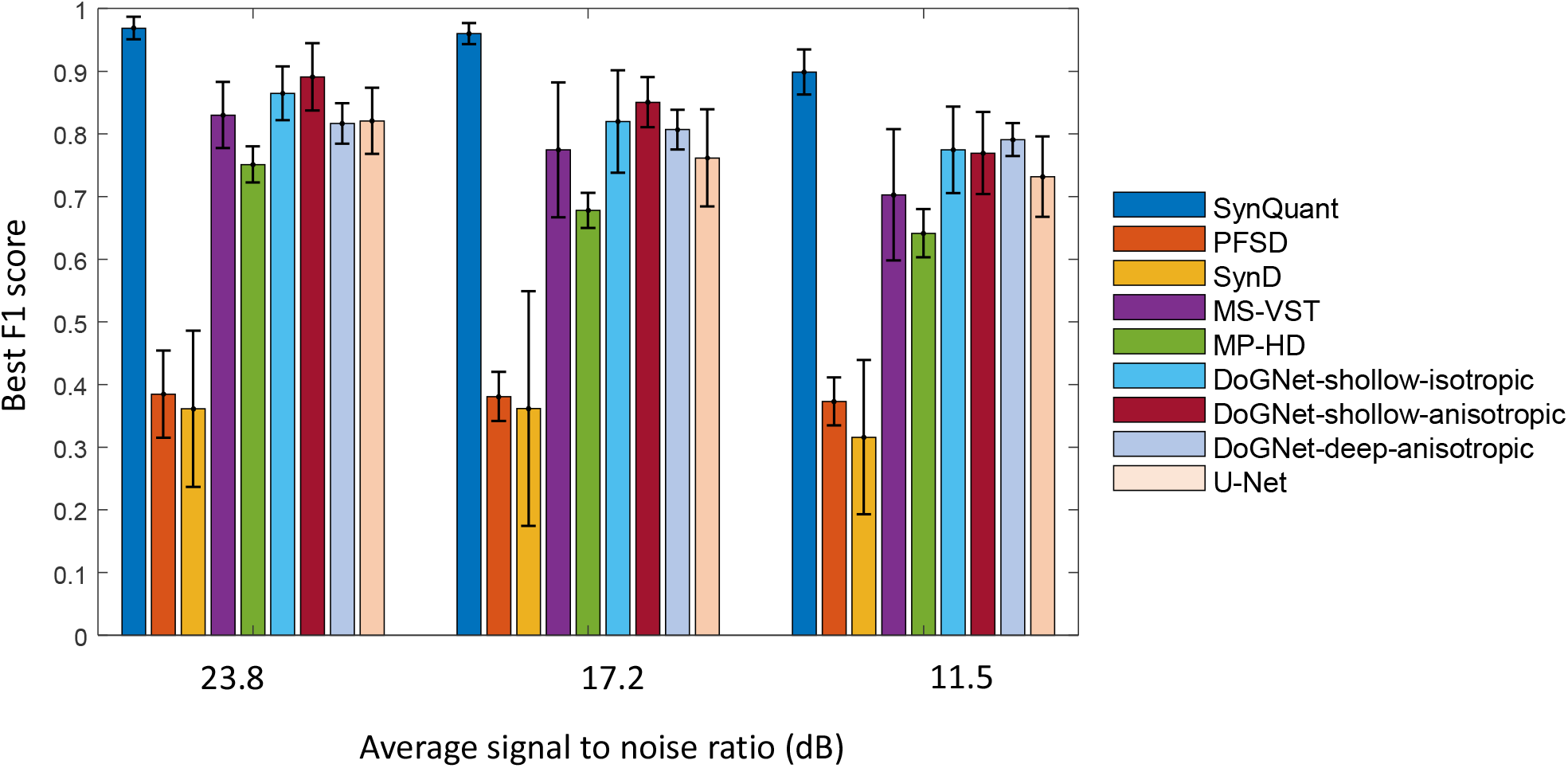
Performance on simulation data with different signal to noise ratios. The range of puncta size is from 9 to 50. The IoU threshold is 0.5. The error bars represent 95% confidence interval.

**Supplemental Figure 6.**
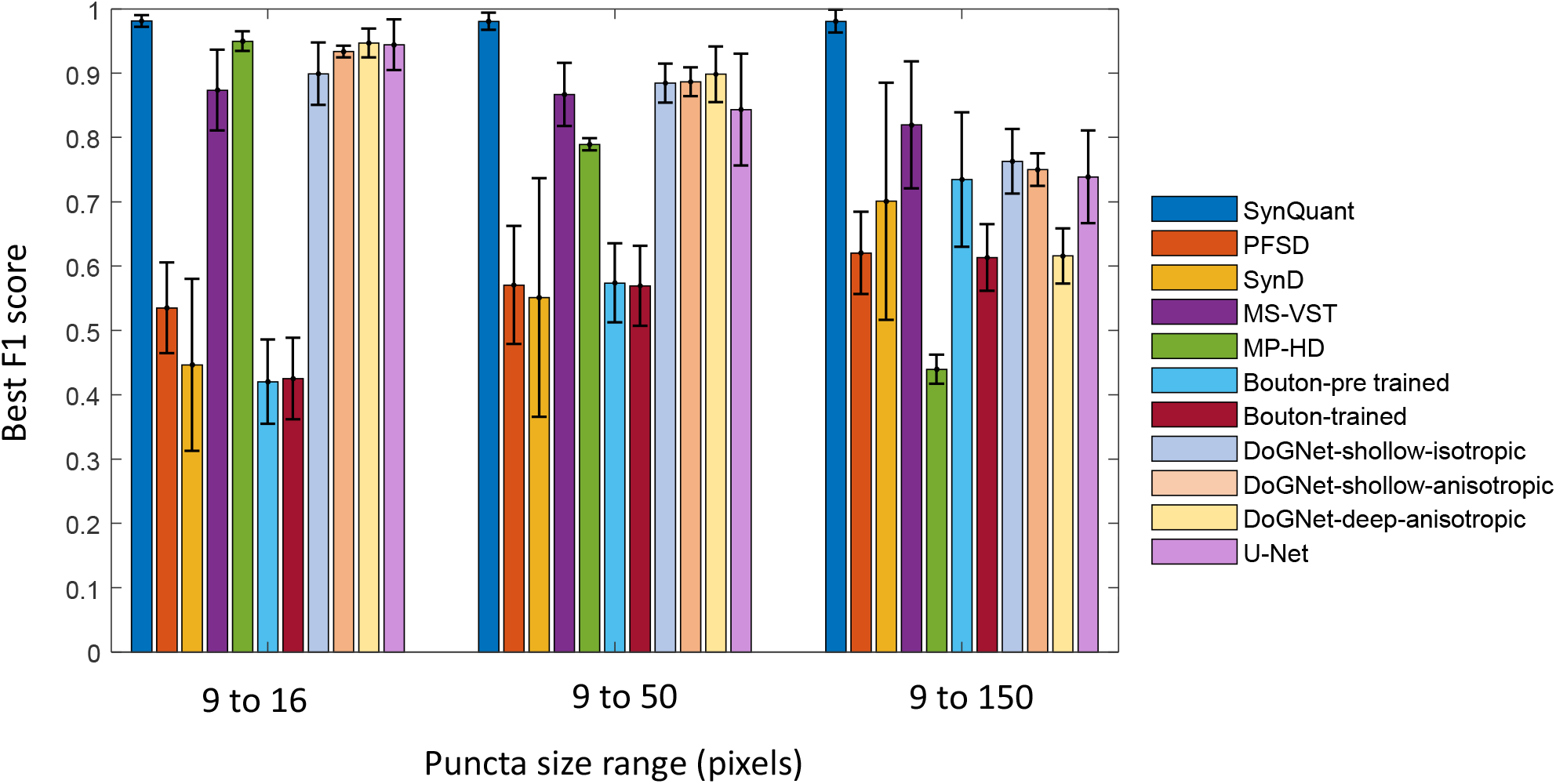
Performance on simulation data with different signal to noise ratios. The SNR is 23.8 dB and the IoU threshold is 0. The error bars represent 95% confidence interval.

**Supplemental Figure 7.**
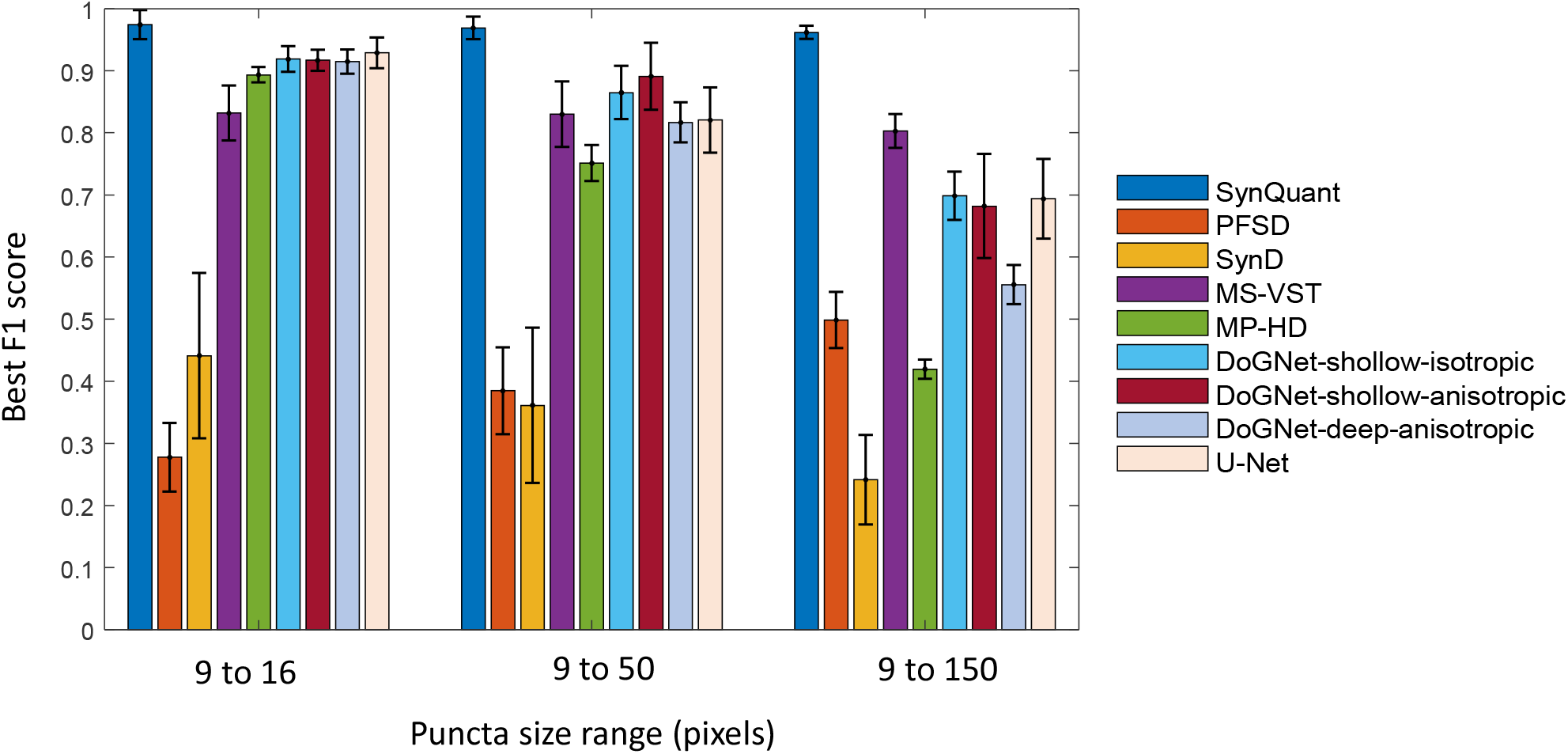
Performance on simulation data with different signal to noise ratios. The SNR is 23.8 dB and the IoU threshold is 0.5. The error bars represent 95% confidence interval.

**Supplemental Figure 8.**
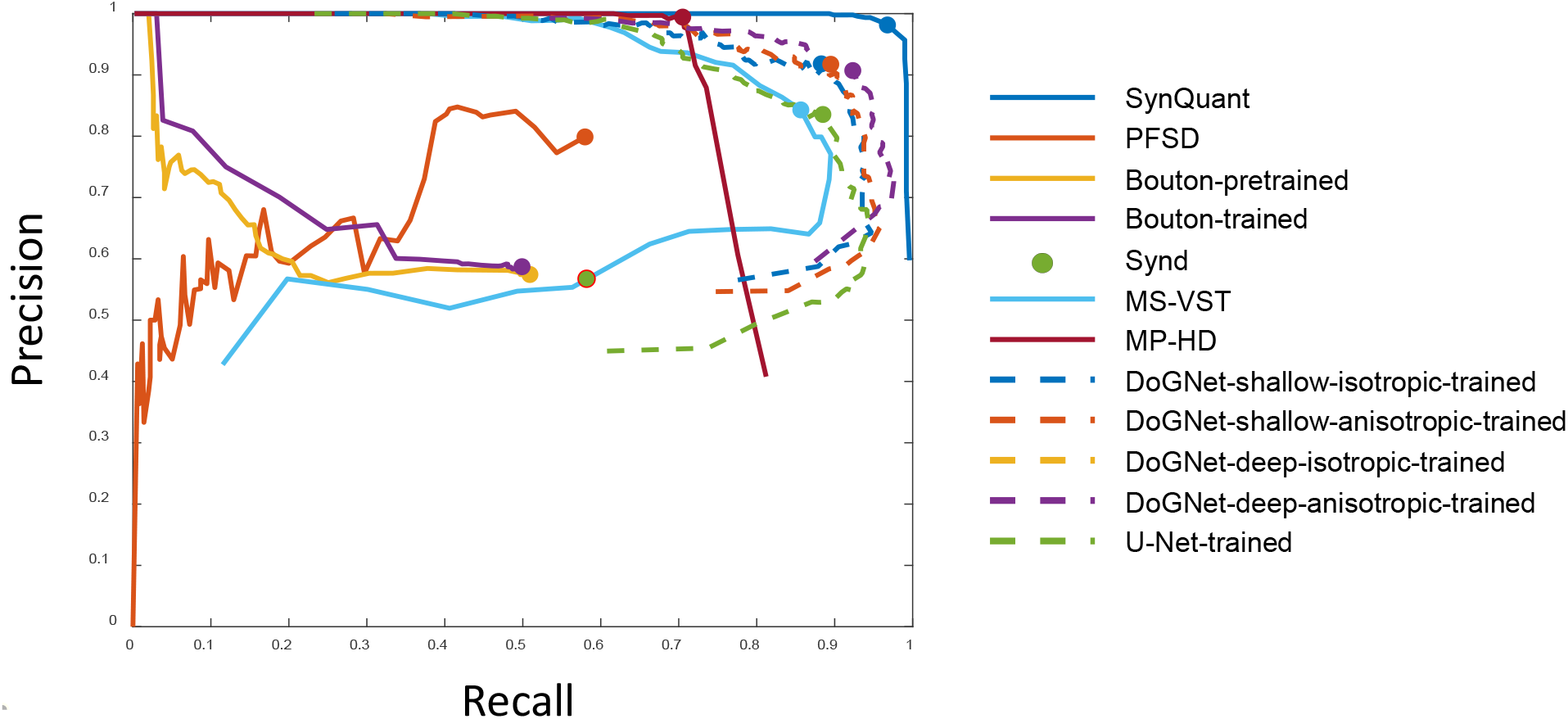
Precision-recall curve for simulation where the puncta size range is from 9 to 50 and the SNR is 17.2 dB. The IoU threshold is 0.

**Supplemental Figure 9.**
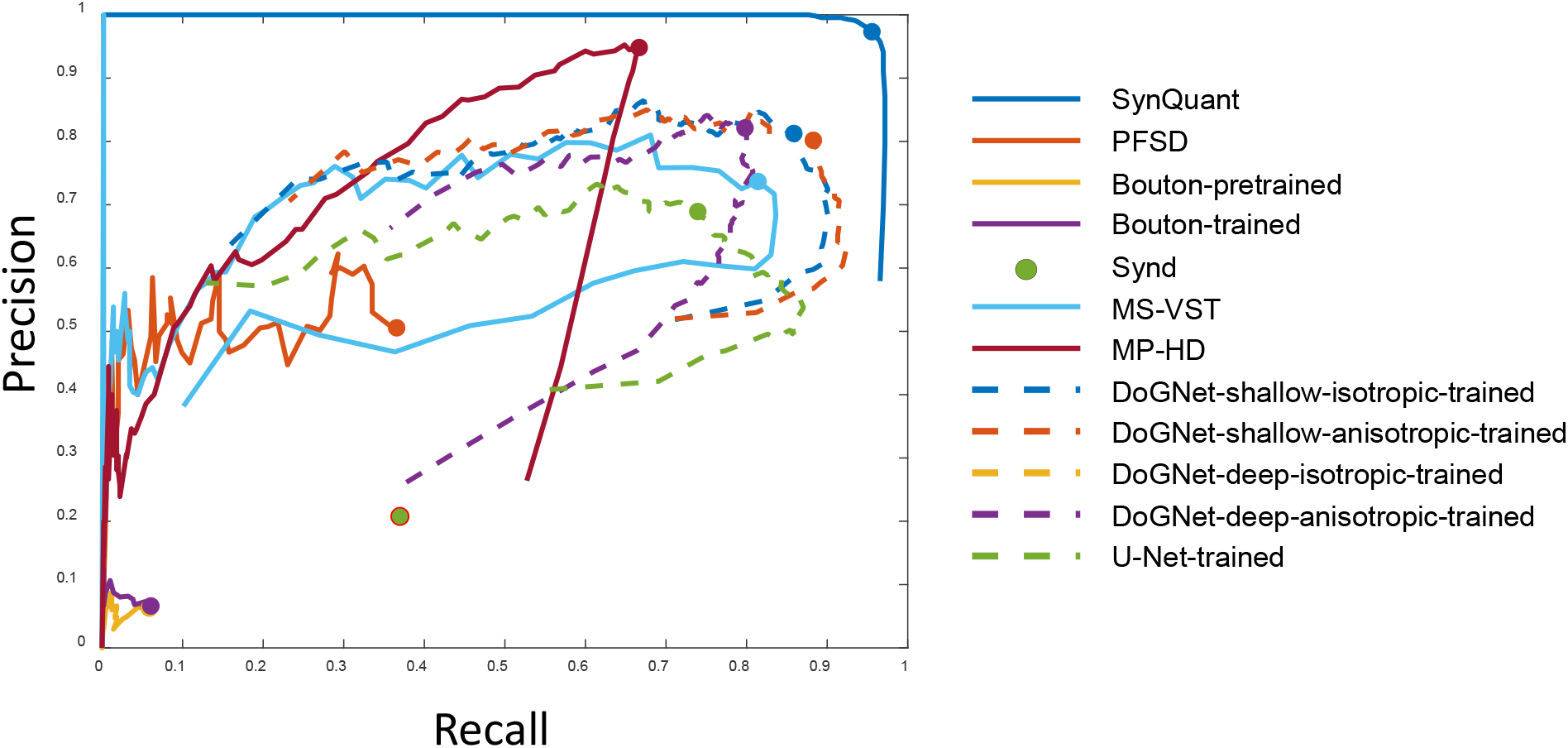
Precision-recall curve for simulation where the puncta size range is from 9 to 50 and the SNR is 17.2 dB. The IoU threshold is 0.5.

### 5 More results on Collman’s data

The ground truth synaptic cleft in EM channel are used in training. However, Bouton needs bounding box around each punctum as the training label Therefore, we only show results with ‘Bouton, pre-trained model’ and do not show results for ‘Bouton, trained’. The results on pre-synaptic channel alone are given in Supplemental Table 2 and the results for post-synaptic channel are given in Supplemental Table 3. In both cases, SynQuant outperforms peer methods in terms of F1 score and average precision. As the post-synaptic channel co-localize well, the final performance of combing two channels is more related to the performance of that channel. In this data set, SynQuant and DoGNet perform the best.

**Supplemental Table 2.** Performance on the pre-synaptic channel in Collman 15 data

| Method | Precision | Recall | Best F1 | Average precision |
| --- | --- | --- | --- | --- |
| **Unsupervised** |  |  |  |  |
| SynQuant (proposed) | 0.579 | 0.504 | **0.539** | **0.436** |
| PFSD | 0.538 | 0.352 | 0.425 | 0.216 |
| MSVST | 0.424 | 0.521 | 0.474 | 0.286 |
| MP-HD | 0.424 | 0.521 | 0.468 | 0.286 |
| **Supervised** |  |  |  |  |
| Bouton, pre-trained model | 0.215 | 0.373 | 0.273 | 0.127 |
| Bouton, trained | N/A | N/A | N/A | N/A |
| DoGNet, shallow, isotropic, trained | 0.506 | 0.339 | **0.406** | **0.170** |
| DoGNet, shallow, anisotropic, trained | 0.540 | 0.305 | 0.390 | 0.167 |
| DoGNet, deep, isotropic, trained | N/A | N/A | N/A | N/A |
| DoGNet, deep, anisotropic, trained | N/A | N/A | N/A | N/A |
| U-Net, trained | 0.267 | 0.254 | 0.260 | 0.067 |

**Supplemental Table 3.** Performance on the post-synaptic channel in Collman 15 data

| Method | Precision | Recall | Best F1 | Average precision |
| --- | --- | --- | --- | --- |
| **Unsupervised** |  |  |  |  |
| SynQuant (proposed) | 0.882 | 0.699 | **0.780** | **0.754** |
| PFSD | 0.885 | 0.589 | 0.707 | 0.666 |
| MSVST | 0.905 | 0.648 | 0.756 | 0.715 |
| MP-HD | 0.876 | 0.648 | 0.745 | 0.680 |
| **Supervised** |  |  |  |  |
| Bouton, pre-trained model | 0.600 | 0.216 | 0.318 | 0.227 |
| Bouton, trained | N/A | N/A | N/A | N/A |
| DoGNet, shallow, isotropic, trained | 0.896 | 0.644 | 0.749 | 0.638 |
| DoGNet, shallow, anisotropic, trained | 0.865 | 0.665 | **0.752** | 0.647 |
| DoGNet, deep, isotropic, trained | 0.820 | 0.686 | 0.747 | 0.672 |
| DoGNet, deep, anisotropic, trained | 0.767 | 0.674 | 0.717 | 0.637 |
| U-Net, trained | 0.849 | 0.669 | 0.749 | **0.680** |

### 6 Examples of real data sets and detected puncta

We show examples of raw data, labels and detection results from all methods for the real and synthetic data sets used in the experiments. For each dataset, we choose one image in the test set to illustrate the detection results. For each method, we find the best F1 score on this set and get the corresponding threshold. Then we apply the threshold on the score map and perform the post-processing steps as discussed in the main body. The IoU threshold is 0 for real data and 0.5 for synthetic data. In general, unless a method performs much worse than the others, it is not easy to assess the performance by eye through these examples.

#### 6.1 Bass’s in vivo 3D data

We choose an image from the test data set in Bass’s data and the results are shown in Supplemental Figure 10. The image is mean projected on the z axis. The label provided by the data set is shown in the third column of the first row. The size of those labels is much larger than the ground truth puncta and we replaced all of them with a 5 by 5 pixels square, as shown in the second column of the first row. When the deep models in DoGNet are trained using the 10 training images in this data set alone, they cannot make correct predictions (fourth row of Supplemental Figure 10). If we first train these deep models with Collman 15 data and tune them with the training data from Bass’s data, they can make good predictions (fifth row of Supplemental Figure 10).

#### 6.2 Neuron astrocyte co-culture data

We show the results of different methods on an image of the neuron astrocyte co-cultured data set in Supplemental Figure 11. Due to the high noise level, some puncta have low contrast with their background. They are removed if they cannot reach consensus among annotators. Again, the deep models of DoGNet cannot be trained by the training data alone. If we tuned it after training with Collman 15 data set, these deep models can make reasonably good predictions.

#### 6.3 Collman’s array tomography data

We show the results on a z stack in Collman 15 data in Supplemental Figure 12. We show one pre-synaptic channel and one post-synaptic channel. We predict the location of synaptic cleft annotated in the electron microscopy channel using these two channels. Note that the post-synaptic channel co-localizes better than the pre-synaptic channel. Besides, some regions with both pre-synaptic and post-synaptic staining do not correspond to the synaptic cleft shown in red. The Bouton method, either pre-trained with the in vivo data or trained with the array tomography data, cannot correctly predict the labels. This might be because the puncta in this data are denser than the *in vivo* data. It is also because the locations of the synaptic clefts are not suitable for training Bouton, which prefers the centers of the fluorescence puncta.

#### 6.4 Synthetic data

We show the results for a synthetic image in Supplemental Figure 13. In that image, the size range of the puncta is from 9 to 50 pixels and the SNR is 17.2 dB. SynQuant performs better than other methods on this data. Interestingly, the deep isotropic model cannot be well trained. Despite that, other models in DoGNet perform well.

**Supplemental Figure 10.**
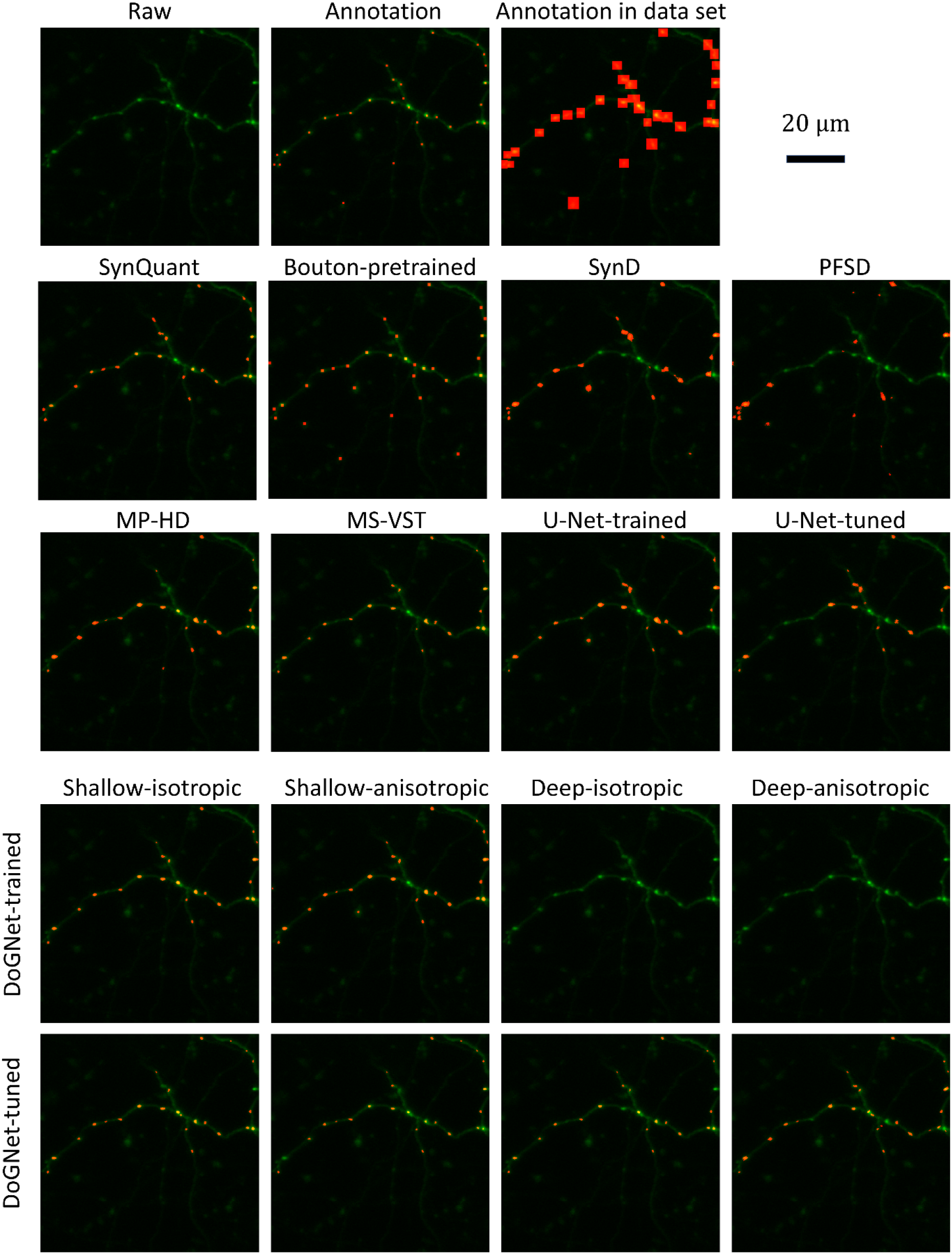
Performance on Bass’s in vivo 3D data. We show the result on one image. The image is mean-projected along the z axis.

**Supplemental Figure 11.**
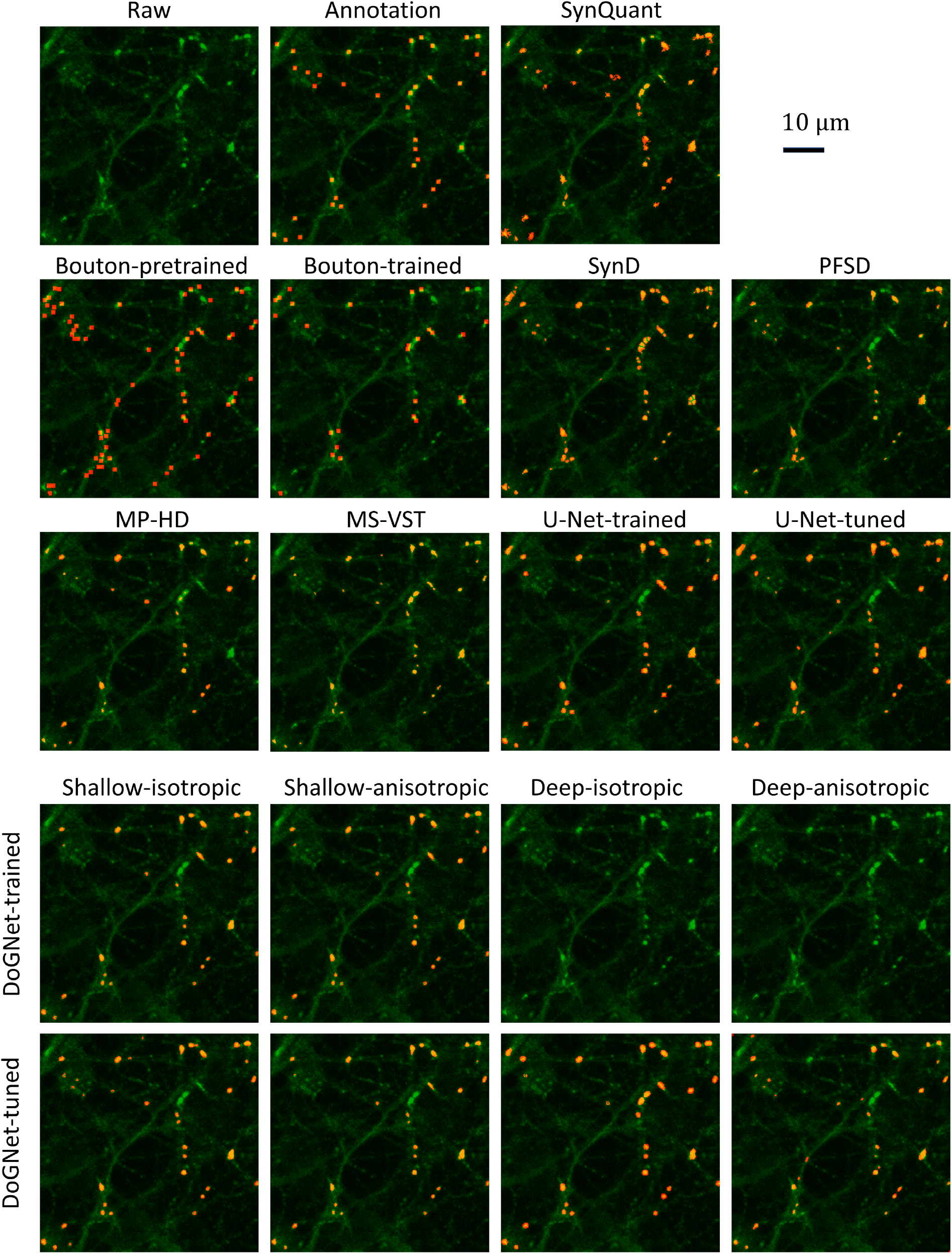
Performance on our in-house neuron astrocyte co-culture data set.

**Supplemental Figure 12.**
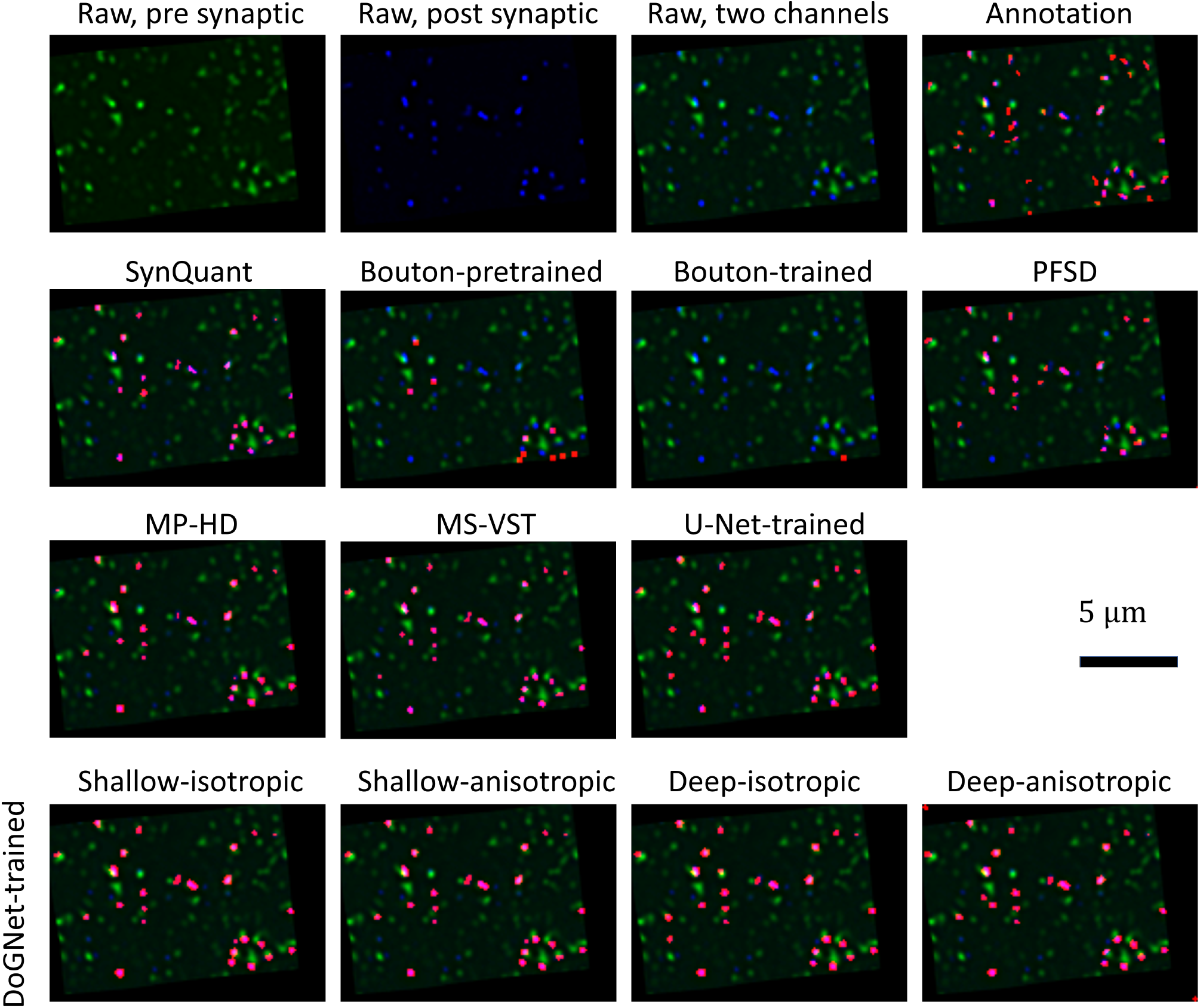
Performance on Collman 15 array tomography data. The green channel is the pre-synaptic channel labeled by Synapsin and the blue one is the post-synaptic channel from PSDr. The red regions are the annotation synaptic cleft on the electron microscopic channel. They are down-sampling to match the resolution of the fluorescence staining.

**Supplemental Figure 13.**
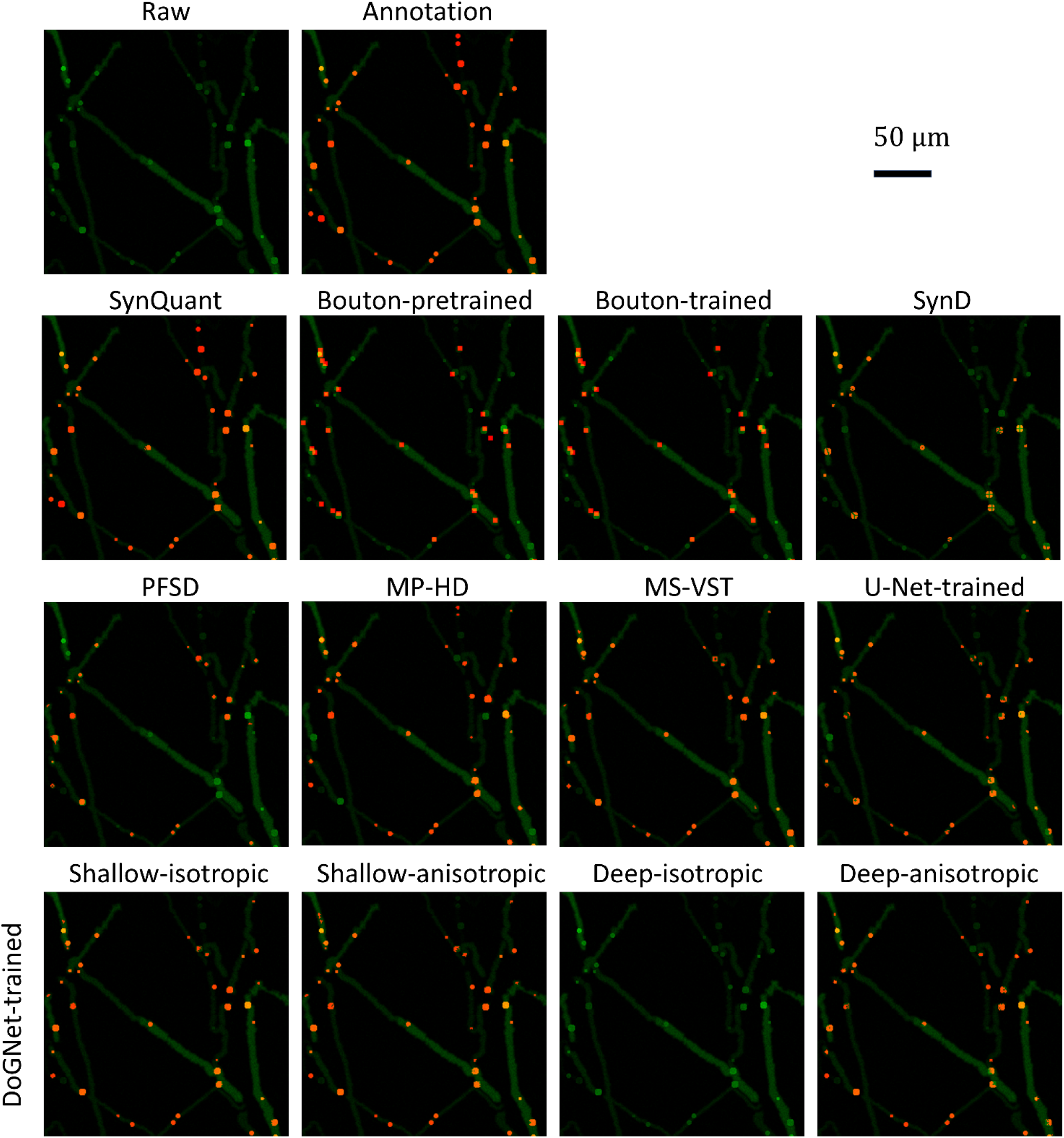
Performance on a synthetic image. The puncta size can range from 9 to 50 and the SNR is 17.2 dB.

### 7 Case study: synaptic puncta and Down-syndrome phenotype

We applied SynQuant to images from a neuron-astrocyte co-culture. An image includes three channels: the nucleus channel (blue) is stained with Hoescht, the pre-synaptic puncta channel (green) is labeled by Synapsin I, and the neurite channel (red) is labeled with Tuj1. We utilized the Synapsin I channel to detect putative pre-synaptic puncta. The red channel was used for neurite tracing and feature extraction.

Synaptic punctum density is defined as the number of puncta per unit length of the neurite. Using punctum density alone will lead to biased analysis results in an inhomogeneous image. For example, in a brighter region, more puncta are expected to be detected, while in the darker region, less puncta can be detected and the density will be lower because of the lower local SNR instead of the true density. This bias is the consequence of region brightness inhomogeneity in the image, which can be fixed by utilizing other neurite and puncta features. In this experiment, we consider the puncta and neurite features on the segment level. From the Synapsin I channel, features of punctum like mean intensity, size, and position are extracted. Neurites are then cut into pieces by their branch points and end points. In one image, we get hundreds of pieces and the intensity of each of them is approximately homogenous inside. Each punctum is assigned to a corresponding neurite piece.

We apply Poisson regression to investigate the relationship between disease status and synaptic punctum density using five groups of samples (Supplemental Table 4). We use the neurite features as confounding factors. DS1 and DS2 are from co-culture samples containing two isogenic Down syndrome (DS) astroglial lines. DS1 astrocytes exhibit trisomy of chromosome 21, the defining hallmark of Down Syndrome. In contrast, the DS2 line, while derived in a manner similar to the DS1 line, instead exhibits disomy of chromosome 21. The DS2 line is thus expected to be more similar genetically to cells derived from non-DS sources. HA is a co-culture sample with human prenatal primary astrocytes. NO are samples from neuron monocultures. The control group is neurons co-cultured with astrocytes differentiated from normal induced pluripotent stem cells (iPSCs). Since we use binary coding for these four groups in the regression model, the control group is the baseline and is not shown in Supplemental Table 4. Length, scale, and intensity are neurite features. First, we only used neurite length as a confounding factor (Supplemental Table 4, column 2 and 3). We found that the NO, HA, and DS2 groups all show significant influence on synaptic punctum density. Then we used all three neurite features as confounding factors (Supplemental Table 4, column 4 and 5). Under these analyses, only HA and NO groups have significant coefficients and DS1 and DS2 do not have an obvious impact on punctum density. From the comparison in Supplemental Table 4, we can see that quantification of punctum density based only on puncta number and neurite length is prone to assigning higher degrees of statistical significance to all the different phenotypes.

The lack of astrocytes in NO group clearly shows the big impact of astrocytes on the formation of synapses (Boehler et al. 2007). Astrocytes in HA group play important roles in synapse formation as well (Reemst et al. 2016, Clarke et al. 2013), but these cells are less mature than the astrocytes in the control group. Therefore, it reduces the synaptic punctum density compared with the control group, but the reduction is less severe than in the NO group. DS2 group is actually similar to non-DS groups, and it is expected to show no significant impact on synaptic punctum density once the confounding factors are controlled. The significant association with neurite length is expected as longer neurite is likely to harbor more synapses, however, the length only cannot account for all neurite heterogeneity. Previous studies have revealed diminished synapse formation and reduced dendritic ramification in infants with DS (Becker et al. 1991; Garner et al. 2012) and lower density of excitatory synapse also were observed in mouse models of DS (Kurt et al. 2004). However, currently, no studies have been able to give a measurement of synaptic density in human brains with DS. As shown here, without proper consideration of confounding factors, biased quantifications can result in misguided conclusions.

Our algorithm does not separate overlapped puncta, because puncta seldom overlap in our in-house data. However, different puncta do have different sizes. It may be unfair to consider puncta with varies sizes equally under such condition. Here we use puncta’s scale rather than the number as a dependent variable for the relationship test. The results are shown in Supplemental Table 5. If we do not consider the confounding factors like neurite scale and intensity, the phenotype DS2 will show a significant effect on the puncta scale.

**Supplemental Table 4.** Poisson regression using Down syndrome phenotypes and three neurite features as predictors and synaptic punctum density as the response

| Features | Neurite length only |  | All neurite features |  |
| --- | --- | --- | --- | --- |
|  | Estimated Coefficients | P-value | Estimated Coefficients | P-value |
| Length | 0.248 | 0 | 0.173 | 1.62e-34 |
| Scale | N/A | N/A | 0.079 | 9.43e-09 |
| Intensity | N/A | N/A | 0.263 | 1.45e-152 |
| DS1 | 0.044 | 0.235 | -0.058 | 0.115 |
| DS2 | 0.156 | **4.43e-06** | -0.060 | 0.086 |
| HA | -0.366 | **1.60e-25** | -0.287 | **3.96e-16** |
| NO | -0.748 | **1.16e-69** | -0.655 | **1.26e-53** |

**Supplemental Table 5.** Poisson regression using Down syndrome phenotypes and three neurite features as predictors and synaptic puncta scale as a response

| Features | Neurite length only |  | All neurite features |  |
| --- | --- | --- | --- | --- |
|  | Estimated Coefficients | P-value | Estimated Coefficients | P-value |
| Length | 1.392 | 0 | 0.596 | 1.77e-12 |
| Scale | N/A | N/A | 0.934 | 1.36e-27 |
| Intensity | N/A | N/A | 1.022 | 3.45e-140 |
| DS1 | 0.210 | 0.072 | -0.136 | 0.241 |
| DS2 | 0.677 | **8.59e-10** | 0.111 | 0.318 |
| HA | -0.916 | **5.95e-21** | -0.651 | **2.55e-11** |
| NO | -1.437 | **7.68e-44** | -1.091 | **4.13e-26** |

### 8 Compare t-test with order statistics with pure noise simulation

We compare Student’s t-test with SynQuant (with order statistics) when evaluating the significance between a region and its neighborhood pixels (Supplemental Table 6). We first simulated an image of size 1024×1024 where all pixels have the same intensity. Then additive Gaussian noises with zeros mean and unit variance were introduced. The threshold varied from 1.2 to 1.8 with step size 0.2. Ideally, no puncta should be detected. First, we assumed the noise variance was known. We counted the total number of regions given by the thresholds and the number of candidates that are significant under the FDR level of 0.05. Totally we got 2574 candidates. SynQuant only reported 12 significant candidates while t-test generated 1310 false positives. So, t-test produces an inflated false positive in thresholding-based methods (Supplemental Table 6). We also observed that the control of the false positives for our method was conservative due to the correlation of scores between overlapped regions. Then we repeated the experiment with a poorly estimated noise variance (0.5 vs true 1.0). Then both tests generated many false positives. This confirms the importance of noise variance estimation in evaluating the significance level for order statistics.

**Supplemental Table 6.** The false positive rate on data with pure noise

| Noise variance used | t-test | Order Statistics |
| --- | --- | --- |
| True noise variance | 0.543 | 0.0047 |
| 0.5 (a poor estimate) | 0.539 | 0.424 |

### 9 Distribution from 1D and 2D null hypothesis

The distribution of the NULL hypothesis in our method is obtained by order statistics. In order statistic theory, if ***x*** is a random vector where each element follows I.I.D. Gaussian distribution, its test statistic (see Equation (2) in the main paper) follows a Gaussian distribution determined by M and N (see Equation (2) in the main paper). Here we test on images, where the signal is in 2D instead of 1D.

We generate an image that contains only Gaussian noises. Then all regions whose intensities are larger than their surrounding pixels are chosen. Based on these puncta, we can draw the distribution of their test statistics. Supplemental Figure 14 shows this distribution, along with the distribution theoretically generated from 1D derivation. For simplicity, here we only show the distribution associated with 30 pixels (15 pixels with higher intensities vs. 15 neighboring pixels with smaller intensities). It’s shown that for images, the order statistics derivation is also applicable.

**Supplemental Figure 14.**
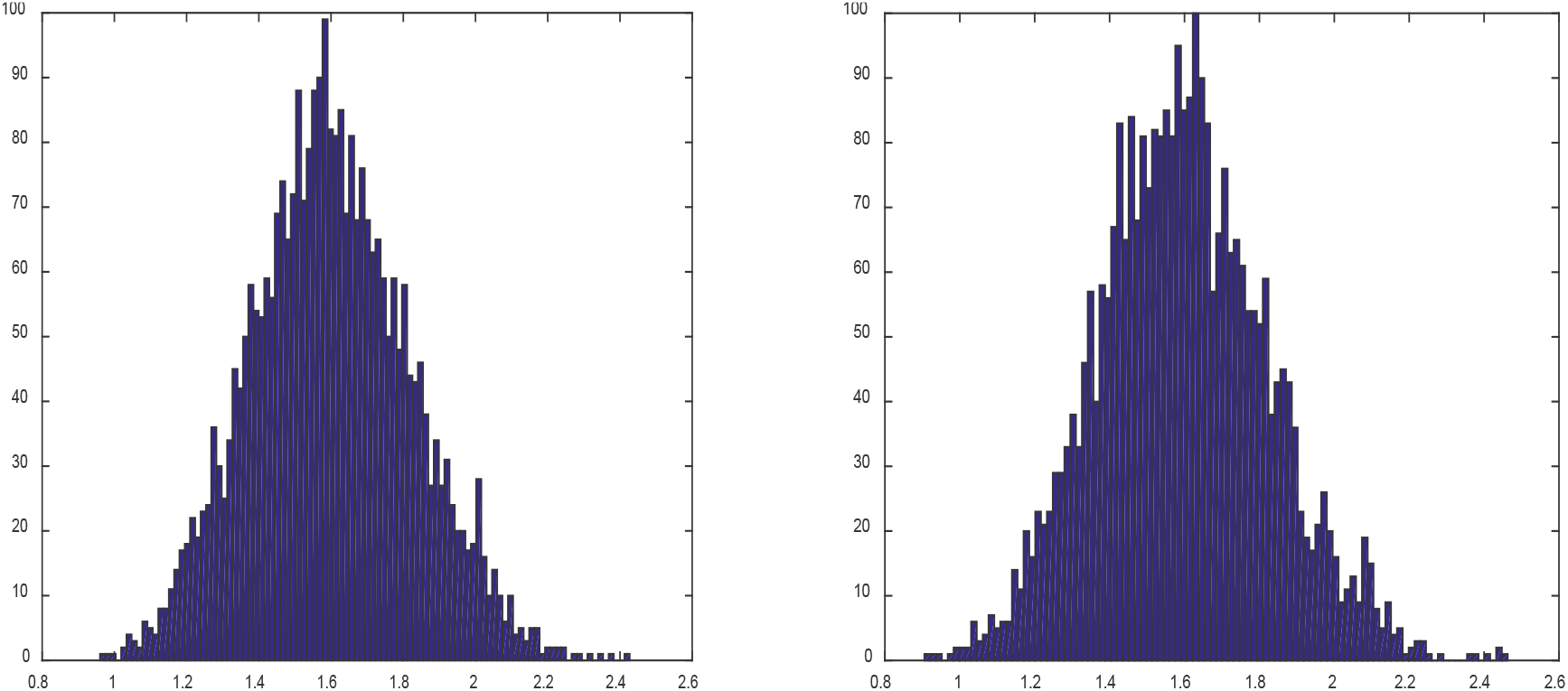
Null distribution on image. The left one is the distribution got from one image with pure noises. The right is distribution got theoretically by generating random vectors. It’s shown that order statistics is also applicable when confronted image data.

### 10 The accuracy of noise variance estimation

The noise of microscopy image intensity consists of two parts: photon noise and camera readout noise. Therefore, it can be modeled by a Mixed-Poisson-Gaussian process (Zhang et al., 2007). The accuracy for the noise variance estimation is essential for the remaining steps in our algorithm. Usually, the Poisson part plays a more important role. Therefore, we first show the noise variance estimation accuracy on our microscopy images.

The model is fitted through the algorithm designed by Foi et al. (2008), though we do not consider the clipping and saturation effects for simplicity and efficiency. The outcome is a linear relationship between mean intensity and noise variance. Ground truth is obtained as follows. The image of one sample in the experiment isf repeatedly taken for ten times under the identical conditions. Each pixel is observed 10 times. Then we can calculate the noise variance for each pixel and plot the relationship between its mean intensity and variance (curve ‘gt’ in Supplemental Figure 15).

From every single image, we estimated the noise and compared them with the ground truth. The comparison is shown in Supplemental Figure 15. It’s clear that the estimated noise model is quite similar to the true value. Note that the linear regression line ‘gts’ in the figure is biased by the pixels with large means and variances. There are a relatively small amount of such pixels and are less reliable. On the contrary, the results from single image estimation (line 1 to 10) are closer to the ground truth (curve ‘gt’).

**Supplemental Figure 15.**
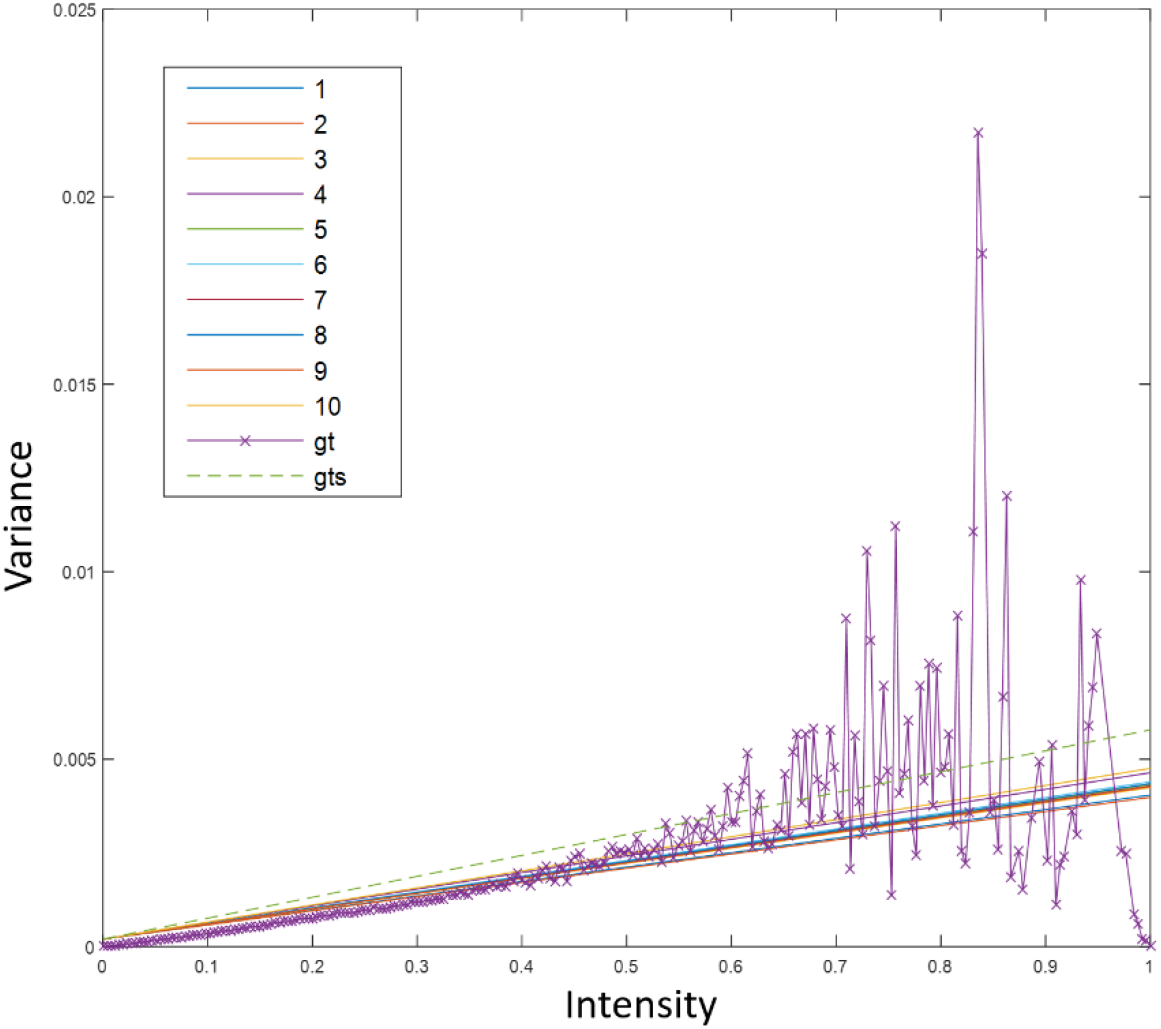
Noise variance estimation results. Line 1-10 are 10 noise models estimated from 10 single images. gt and gts are the ground truth from 10 images and the linear regression result, respectively.

### 11 Blind source separation for neurite-like signal removing

As we had access to both the Synapsin I channel (green channel) and the neurite channel (red channel), we attempted to use the neurite channel to remove the neurite-like background in the Synapsin I channel. This procedure is like blind source separation (BSS). Under the assumption that the neurite channel is mostly pure, whose signal mainly from true neurite, the Synapsin I channel can be viewed as a mixed signal from the known neurite channel and unknown pure synaptic puncta signal. If we could extract the pure synaptic puncta signal, the difficulty of synaptic punctum detection and segmentation problem should be decreased.

For any two pixels *i* and *j*, the intensities in neurite channels are *y*_*i*_ and *y*_*j*_. The intensities in Synapsin I channels are *x*_*i*_ and *x*_*j*_. If we assume that signals in neurite channel go into the Synapsin I channel, we will have *y*_*i*_ = *a*_*i*_*x*_*i*_ + *b*_*i*_ and *y*_*j*_ = *a*_*j*_*x*_*j*_ + *b*_*j*_. In order for blind source separation to work, we require *a*_*i*_ = *a*_*j*_ and *b*_*i*_ = *b*_*j*_. If this is true for any pixels, we say the relationship between these two channels are homogeneous. Otherwise, if *a*_*i*_ ≠ *a*_*j*_ for different pixels, the relationship is inhomogeneous and the blind source separation is not a valid choice.

Here we first show the inhomogeneous relationships using the real data. Then we list several BSS attempts based on regression and non-negative matrix factorization (NMF). Both the analysis and experiment results show that the Synapsin I channel is not simply a linear combination of synaptic puncta and neurite signals and thus making the separation difficult to achieve.

#### 11.1 Inhomogeneity of relationships between two channels

If *y* is the intensity of one pixel in the Synapsin I channel, and the *x* is the intensity for the same pixel in the neurite channel, their relationship is given by a regression model

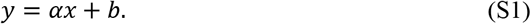

Here *α* is a random variable. We assume it follows Gaussian distribution *N*(*μ*, *σ*^2^).

To study whether the relationship between these two channels is homogenous, we cut the neurite into small (homogeneous) pieces. The idea is that we study the relationship piece by piece. If the relationship is homogeneous, the results from difference neurite pieces should be similar.

Assume we have *M* pieces. Also, assume *y*_*i*_ is the intensity of any pixel in piece *i* in Synapsin I channel and *x*_*i*_ is the intensity for the same pixel in the neurite channel. We have a new regression model

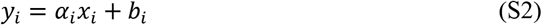

For piece *i*, *α*_*i*_ consists of three components

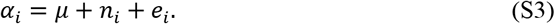

Here *e*_*i*_ is the noise caused by sample size (pixel number) in piece *i* and *n*_*i*_ is the term related to the position or other factors. Consider all pieces, we have the regression model for coefficient *α*

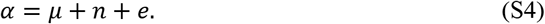

If the relationship between these two channels are homogeneous, *n* is zero and thus all *α* are determined by *μ* and *e*

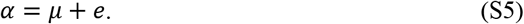

Under the homogeneous condition, across piece variance *var*(*α*) should be equal to within-piece variance *var*(*e*), otherwise, *var*(*n*) is n on-zero. By calculating *var*(*α*) and *var*(*e*) from real data, we can find whether the relationship between these two channels are homogeneous.

We stabilize Synapsin I and neurite channels to make the noise variance un-related to intensity. Then in neurite channel, we cut neurite into small homogeneous pieces and remove those whose skeleton is less than 10 pixels. The pixels inside one piece in these two channels should be linearly related as is illustrated in Equation (S2). For each *α*_*i*_, 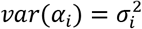 is got from the linear regression. Taking the average of 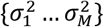, we obtain *var*(*e*). With {*α*_1_, …, *α*_*M*_} we can estimate the sample variance *var*(*α*).

Using the two channels shown in Supplemental Figure 1(B), we get that *var*(*α*)=0.064 and *var*(*e*)=0.015, which means *var*(*n*) is not zero (0.049) and the relationship between these two channels are inhomogeneous. By Chi-Square test, the *p*-value of *var*(*α*)=0.015 is 0, which means it’s almost impossible to happen.

#### 11.2 Linear regression

Here we test the relationship of neurite in these two channels. To reduce the influences of synaptic punctum signals, we remove most puncta before the test. Then linear regression is applied to the remaining pixels in the two channels. We subtract neurite channel based on the regression results, as shown in the bottom two figures of Supplemental Figure 16. Though it seems most neurite-like signals have been removed from observation, the remaining artifacts introduce more interference.

#### 11.3 Non-negative matrix factorization (NMF)

NMF is a famous blind source separation technique. For our problem, we assume the channels are both composed of pure synaptic puncta and neurite signals. To apply NMF, we use the Synapsin I and the neurite channels to form a 2 by *N* matrix V, where *N* is the number of pixels in each channel. In practice, for each channel, all horizontal rows of pixels are concatenated into a 1-D (column) vector and then be transformed into a 1-D (row) vector. These two vectors form the 2 by *N* matrix V. By NMF, V is factorized into a 2 by 2 matrix W and 2 by *N* matrix H. The two vectors in H should represent pure synaptic puncta and neurite signals. Results of NMF are shown in the right-up figure in Supplemental Figure 16. Because *N* ≫ 2, the un-mixing results are un-reliable. It’s obvious that the synaptic punctum signals extracted from NMF are as noisy as before.

**Supplemental Figure 16.**
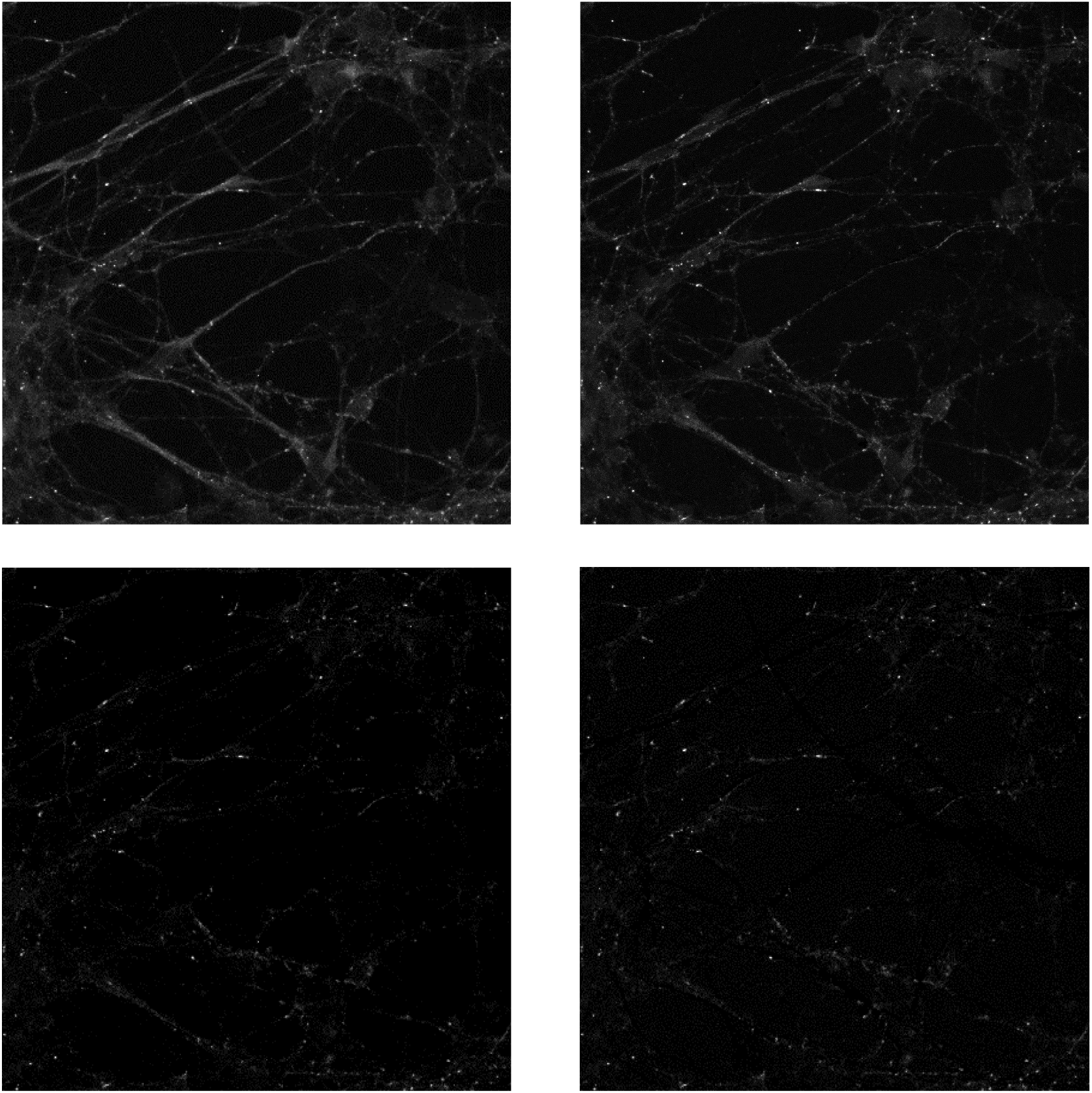
Results of BSS trials. The top-left is the original synapse channel. Top-right is the result got from NMF. It’s obvious that NMF fails to handle most of the neurite-like signals. The bottom-left and bottom-right are results of linear regression with and without bias term. These two results are similar and looks like that most neurite-like signal has been removed. Our following experiments shows that this process introduces un-seen artifacts which does not truly help for detection.

### 12 FDR control

The FDR control offers a looser restriction over Bonferroni correction. We conduct FDR control in each iteration by testing whether all the un-selected nodes (regions) fail to meet the significance request. Otherwise, we pick up the most significant remaining one and continue to the next iteration until no such region exists. For example, as shown in Figure 1(H) in the main paper, we have two candidate synaptic puncta **b** and **f**. The next node with the highest significance value is **c**. Then FDR control will be conducted on all the nodes including **b** and **f** to see under such condition whether **c** is still significant. If so, set **c** as a candidate punctum. Otherwise, stop and report all candidate synaptic puncta we detected. Because for each iteration, the significance (order statistics score) of some nodes may change, FDR control is needed in each iteration. As overlapped regions may be highly correlated, for FDR control, we use the general case introduced by Benjamini and Yosef, (1995).

### 13 Neurite segmentation

For synaptic puncta quantification, it is important to collect puncta properties along with features from neurites. However, the neurite related features obtained from existing tools are not sufficient. For example, existing synaptic puncta quantification tools only utilize the length of neurite, with no other confounding factors in calculating synaptic punctum density. However, this density could be influenced by spatial inhomogeneous factors such as neurite scale or sizes (Glynn et al., 2011; Klintsova and Greenough, 1999). When we test the relationship between synaptic punctum density and related disease phenotypes, if we do not consider such space-related confounding factors of neurites, the conclusion might be unreliable. Therefore, analyzing the properties of neurites in detail will help the study of the physiology and pathology of synapses.

Similar to SynD (Schmitz et al. 2011), we use a steerable filter (Meijering et al. 2004) to trace the neurite. The filter is based on the second-order derivatives of a Gaussian kernel:

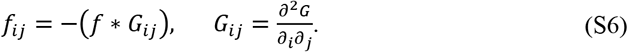

Here * denotes convolution and index i and j are either x or y directions and f is the image. For each pixel, we get the Hessian matrix

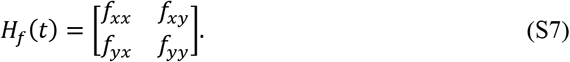

Here *f*_*xy*_ = *f*_*yx*_. The probability that pixel *t* belongs to a neurite is obtained from the eigenvalues and eigenvectors of this matrix. The cost for pixel q to be added to a neurite where pixel p is already on it is

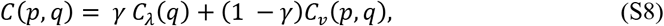

where *γ* ∈ [0,1] is a user-defined weight. In default we set it to be 0.5, so we have no preference for each term. *C*_λ_ (*q*) and *C*_*v*_(*p, q*) are two normalized cost components. The former one is the cost of pixel q to be added to **any** neurite. Here

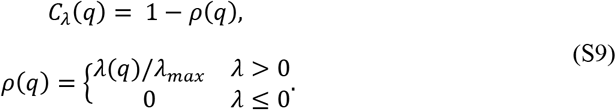

λ_*max*_ is the maximum among the largest eigenvalues obtained from the Hessian matrices of all pixels. λ(*q*) is the largest eigenvalue of pixel q’s hessian matrix. The latter one is the direction cost if we add pixel q to a neurite where pixel p is on it. It is defined as

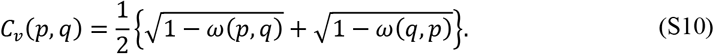

*ω*(*p,q*) = |*v*(*p*)· *d*(*p,q*)|, which is the unit “link vector” from pixel p to q. *d*(*p,q*) = (*q* − *p*)/||*q* − *p*|| and v(q) is the normalized eigen vector of p. By taking the absolute value, this cost component ignores direction and only consider orientation similarity.

With Equation (S8), we can extract neurites from seed points by measuring if the cost is less than a threshold (0.9 in our implementation). Although neurites must grow from somas in the SynD algorithm, this restriction is relaxed in our experiments to avoid false negatives caused by missing somas. In our experiments, every unchecked point would be tested as a seed to find whether it belongs to a nearby neurite.

The detected neurite is cut by branch points and end points for further analysis. We assume the intensities of pixels in each piece is similar. Each piece corresponds to one or more synaptic puncta that stay on it. Concerning both synaptic punctum features and related neurite features, a comprehensive quantification of the synaptic puncta can be conducted.

An example of neurite tracing result is given in Supplemental Figure 17.

**Supplemental Figure 17.**
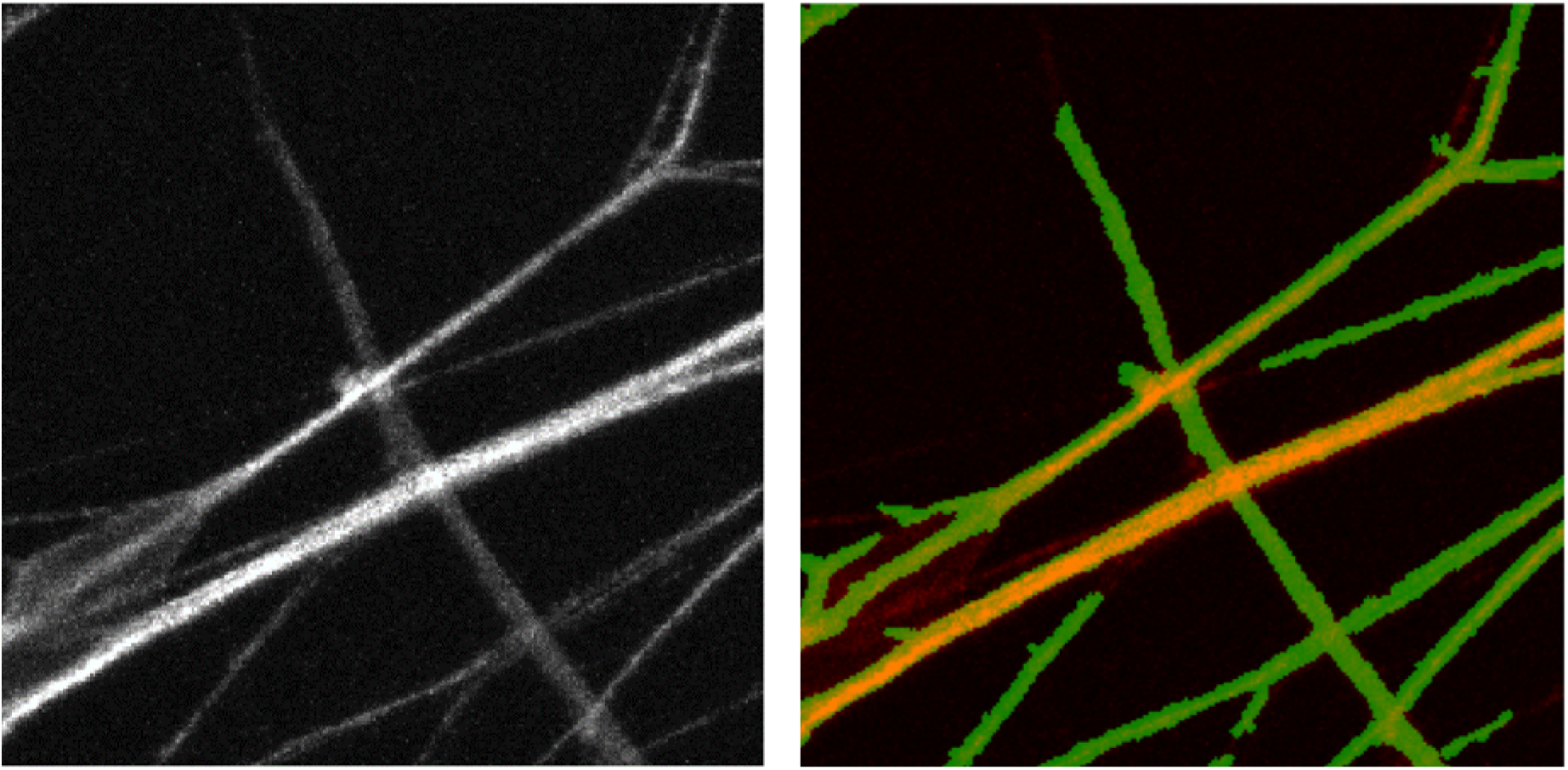
Neurite tracing results. Left: raw data in the Tuj1 channel which stain the neurites. Right: detection results shown in green, which is overlaid with the red Tuj1 channel.

### 14 Implementation and speed

Since the features (used in Eq. 6 in the main text) for *p*-value calculation are accumulative, the tree structure shown in the lower part of Fig.1 in the main text can be built efficiently with adaptive local thresholding (Ranefall et al. 2016). Updating the nodes’ order statistics scores consumes most of the time. Generally, SynQuant takes 1 to 20 seconds to detect and segment synapses on images we tested in the experiments. The speed is influenced by the size of image, depth of image, puncta size range and puncta number. All experiments are performed on a workstation with Xeon E5-2630 CPU.

